# Promoter-responsiveness to distal regulatory elements is mediated by the promoter-proximal region

**DOI:** 10.1101/2025.10.14.682013

**Authors:** Blanka Majchrzycka, Miles K. Huseyin, Ekin D. Aksu, Anders S. Hansen, Stefan Mundlos, Arnaud R. Krebs, Daniel M. Ibrahim

## Abstract

Gene promoters induce transcription in response to distal enhancers located in large gene regulatory domains. How this long-range regulation specifically activates the target promoter while bypassing other promoters located in the same regulatory domain remains unclear. Here, we find that the promoter-proximal region is critical for cell-type specific promoter responsiveness. Using high-throughput genome-engineering in mouse embryonic stem cells (mESC), we systematically replace the endogenous *Sox9* promoter with different libraries of core and extended (i.e. full) promoters and assess their response to long-range regulatory elements in mESCs and neural progenitor cells. We find that only a subset of full promoters is activated by distal neuronal enhancers and that the promoter-proximal region is necessary for this promoter responsiveness. Core promoters alone are insufficient to respond to distal enhancers but modulate the transcriptional output of responsive promoters. Our results suggest that within multipartite regulatory domains, the promoter-proximal region fulfills a facilitator-like function that filters and transmits signal from distal enhancers, ultimately conferring promoter responsiveness to long-range regulatory elements.

**Highlights:**

1. Promoter-RMCE at the *Sox9* regulatory domain demonstrates locus-specific promoter responsiveness in mammals
2. The promoter-proximal region determines and is necessary for long-range enhancer compatibility
3. Core promoters alone are insufficient for responsiveness to distal enhancers
4. The promoter-proximal region has a facilitator-like role as a selective filter within cis-regulatory domains

## Introduction

Cell type-specific gene expression is controlled by the combinatorial action of gene promoters and distal enhancers. Usually multiple enhancers cooperatively induce transcription at their target promoter while skipping genes located more proximal to the enhancers ^1^. Topologically associating domains (TADs) restrict the possible targets of enhancers and often reflect the regulatory landscape of that gene ^2^. However, the same TAD typically contains multiple genes with differential expression patterns ^3–5^. Therefore, a key unresolved question in the field remains: How do enhancers specifically activate one promoter without affecting others?

Various lines of evidence suggest some degree of enhancer-promoter compatibility that helps enhancers to activate specific promoters. In *Drosophila*, housekeeping and developmental core promoters respond very differently to the same enhancer. This was first discovered through detailed analysis at individual loci ^6–8^ and later substantiated through massive parallel reporter assays (MPRA) ^9^. However, whether similar rules apply in mammals remains unclear. Systematic screens using MPRAs have yielded conflicting results ^10–12^. Cell type-specific looping of enhancers to distinct promoters, for example mediated by CTCF-sites ^13–15^ or orphan CpG islands next to enhancers ^16^, can explain certain cases of selective promoter activation. However, such looping mechanisms do not explain the majority of differential promoter responses^17,18^. One example are the *MyfC* and *Myf5* genes, which are separated by only 8kb and show differential responses to a nearby enhancer during myogenic differentiation^19^. Another conundrum comes from pathogenic structural variations in cancer and congenital disease^20^. These can cause gene mis-expression by rearranging enhancers with a new set of promoters ^20–25^. However, in many cases such rearrangements lead to mis-expression of only a subset of genes, while others remain unaffected^23,24,26^. Why some genes respond to such rearrangements while others do not remains hard to predict.

Several technical aspects limit our current understanding of enhancer-promoter compatibility. Traditional reporter assays as performed in many MPRAs lack the genomic context. Episomal and genome-integrated enhancer-reporter assays usually combine relatively short (200-350bp) enhancer and promoter sequences in direct vicinity. In contrast, most mammalian regulatory domains contain multiple enhancers and promoters at various distances. Locus-specific genome-engineering is better suited to study the complexity of endogenous gene regulation, but has so far been limited in throughput, hindering systematic comparisons of many transgenes at the same locus. Recent advances in genome engineering have started to overcome this limitation in throughput^27–29^ revealing the limited concordance between episomal MPRAs and locus-engineered assays^28^, highlighted effects of promoter competition^27^, or demonstrated diminishing promoter activation with increasing enhancer distance^29–32^. The synergistic and hierarchical relationships between *cis*-regulatory elements (CREs) within multipartite regulatory domains have started to be addressed by creating synthetic regulatory domains^31,33–36^. Recently, some of these studies proposed a novel class of CREs – facilitators – which function only as weak enhancers in isolation, but boost the activity of a stronger enhancer when placed between the enhancer and the target gene^31,33,37,38^. Another important feature is the specific position of the promoter within a regulatory domain, which is optimized for transcriptional activation^39,40^. However, which specific role the promoter sequence might play in a multi-enhancer regulatory domain remains unclear.

Importantly, gene promoters can and have been defined in different ways. Best established and understood is the core promoter (~100bp surrounding the Transcription Start Site, TSS) as the assembly platform for the pre-initiation complex^41^ and these show differential enhancer compatibility in *Drosophila*^41–43^. However, one prominent feature of promoters is the promoter-proximal region directly upstream of the core promoter, sometimes referred to as the proximal enhancer^44^. These sequences resemble enhancers in length (~1-2kb) and transcription factor binding site (TFBS) content and are marked by promoter-associated histone modifications. We define the combination of the promoter-proximal region and the core promoter as the full promoter. Full promoters of housekeeping genes (e.g. *PGK*, *EF1a*) are widely used as generic and strong activators in many transgenes and some full promoters of developmental genes are used as tissue-specific drivers in transgenic animals (e.g. *Prr”1* ^45^, *Zp3*^46^). However, most full promoters cannot drive robust, tissue-specific transcription by themselves^47^, posing the question of which other function they might have.

We hypothesize that the promoter-proximal region confers the transcriptional response to distal enhancers in the genomic context of a multipartite gene regulatory domain. We test this using a locus-specific genome-engineering strategy, Promoter-RMCE (Recombinase-Mediated Cassette Exchange). We replace the full promoter of *Sox9* with dozens of promoters from other developmental and housekeeping genes in mouse embryonic stem cells (mESC) and profile their response to distal *Sox9* enhancers during neuronal progenitor cell (NPC) differentiation. Our results show selective response of full promoters to the *Sox9* gene regulatory domain and that the promoter-proximal region is necessary for this responsiveness. In contrast, core promoters alone are insufficient to respond to the same regulatory stimulus but can modulate the transcriptional output of a responsive promoter. We propose a facilitator-like function for the promoter-proximal region of mammalian genes. Together, our results suggest selective enhancer-promoter compatibility in mammals and demonstrate that the promoter-proximal sequences are essential for determining promoter responsiveness to a complex cis-regulatory domain.

## Results

### The *Sox9* promoter-proximal region is essential for activation by distal enhancers

We selected *Sox9* as a model locus to test how different promoters respond to the cell type-specific activity of a large regulatory domain. A multitude of cell-type specific distal enhancers, distributed throughout a ~1.6Mb TAD, activate the *Sox9* gene^26,48–51^. *Sox9* is not expressed in mESCs, but can be induced through *in vitro* differentiation into NPCs (**Fig. S1A**) ^52^. Using ATAC-seq and H3K27ac ChIP-seq in mESC and derived NPCs, we identified 15 NPC-specific and 4 shared mESC/NPC candidate enhancers within the Sox9 TAD (**Fig. 1A, Fig. S1B**). We next performed region capture Micro-C (RCMC)^17^ to assess the 3D chromatin interactions within the TAD in mESCs and NPCs (**Fig. 1A, Fig. S1**). In NPCs, overall interactions within the TAD were increased and virtual 4C from the *Sox9* promoter showed increased interactions with the major CTCF sites^51^ and with NPC-specific enhancers throughout the Sox9 TAD while losing mESC-specific interactions (**Fig. 1A, Fig. S1**). As only protein-coding gene within the TAD, this setup demonstrates that the *Sox9* promoter forms NPC-specific interactions with a multitude of distal CREs, suggesting that they drive *Sox9* induction in during NPC differentiation. The *Sox9* promoter itself has a bivalent chromatin signature surrounding the TSS by several kb in mESCs, which is replaced with active H3K4me3/H3K27ac marks in NPCs (**Fig. 1B**). To test the importance of the promoter-proximal region for *Sox9* expression, we created mESC lines in which we heterozygously deleted either the *Sox9* full promoter (ΔFP) or only the promoter-proximal region (ΔProx) (**Fig. S2A**). Both lines lost 40-50% of *Sox9* expression in NPCs, suggesting full loss of expression on the targeted allele (**Fig. S2B**).

**Figure 1:**
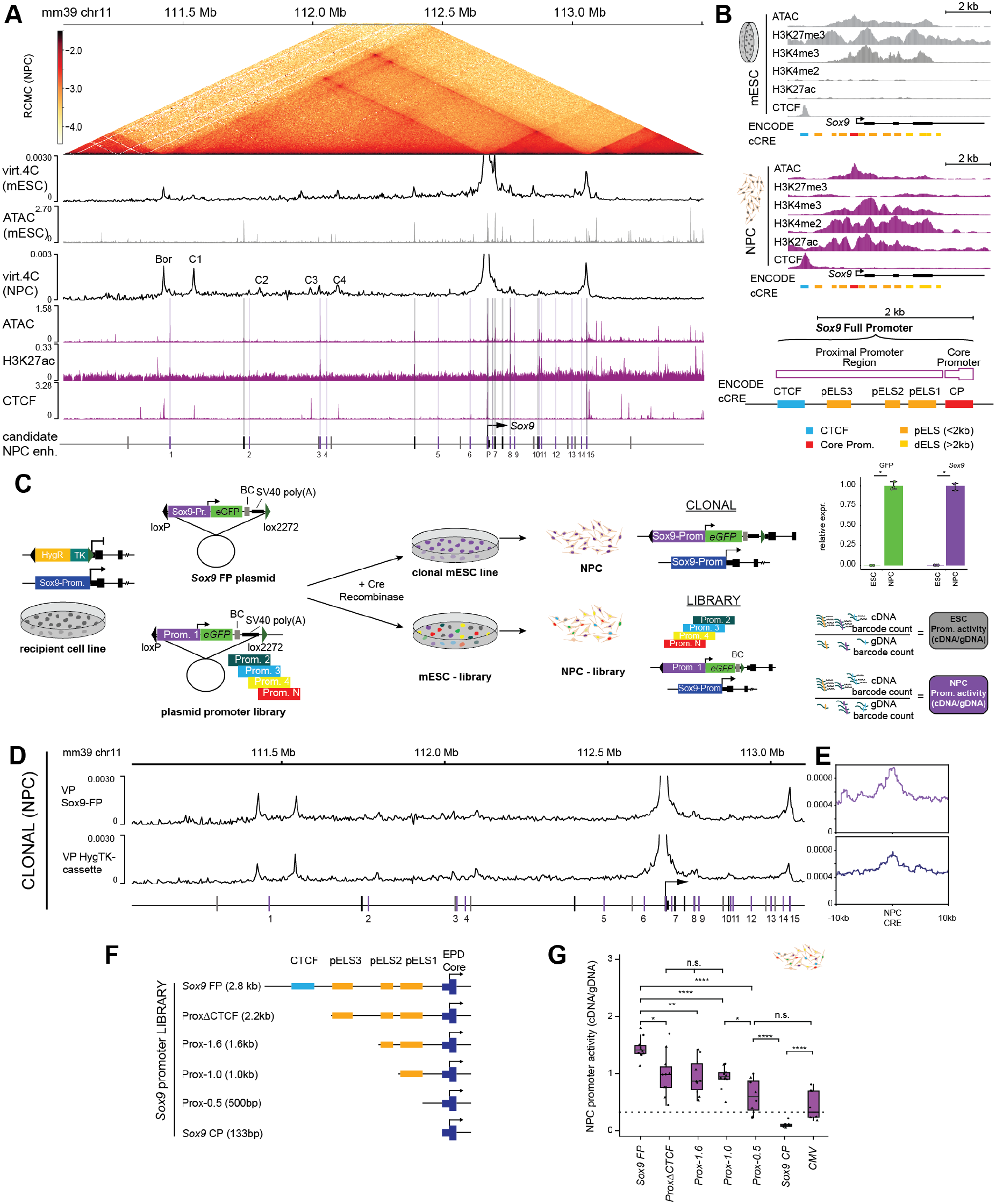
The *Sox9* promoter is regulated by distal regulatory elements during neuronal differentiation. (**A**) RCMC contact maps and virtual 4C profiles from the Sox9 promoter in mESCs and NPCs, alongside ATAC-seq, H3K27ac, and CTCF profiles highlighting candidate NPC enhancers. (**B**) Chromatin signatures at the *Sox9* promoter in mESCs and NPCs. ENCODE classification of promoter-proximal elements and schematic representation of full, core and proximal promoters in this study. pELS: proximal enhancer-like signature, dELS: distal enhancer-like signature (**C**) Schematic of the Promoter-RMCE setup and quantification using RT-qPCR (clonal cell line) and Amplicon-seq (library) (**D**) virtual 4C interaction profiles of NPC cells with Sox9-FP or the HygTK selection cassette viewpoint. Positions of NPC-specific ATAC-seq peaks indicated. (**E**) 1D aggregate RCMC pileups of the respective promoters with the NPC-specific enhancers −/−10kb. (**F**) Schematic representation of variant *Sox9* promoters used in Promoter-RMCE (**G**) NPC Promoter-RMCE activity of variant *Sox9* promoters. Triangles and dots = barcode values from two biological replicates. Dashed line = median CMV activity. p-value < 0.05 from Wilcoxon test: * < 0.05, ** < 0.01, *** < 0.001, **** < 0.0001.)

### Distinct promoter-proximal sequences are necessary for full *Sox9* expression

To systematically explore the role of the promoter in a gene regulatory domain with many long-range enhancers, we established a genome-engineering assay, Promoter-RMCE, designed to replace the endogenous *Sox9* full promoter on one allele with a variety of different promoters. For this, we first inserted a Hyg-TK selection cassette ^53^ flanked by heterotypic *lo”* sites at the deletion breakpoint in the ΔFP mESC line. This generated a recipient *Sox9* Promoter-RMCE mESC line (**Fig. 1C, Fig. S2C**), which allows directed, heterozygous integration of transgenes at the site of the *Sox9* promoter using RMCE (**Fig. 1C, Fig. S2C**). We first tested the system by re-integrating the *Sox9* full promoter driving an *eGFP* transgene, derived a clonal mESC line and measured *GFP* and *Sox9* expression in mESCs and NPCs using qRT-PCR. Both, *Sox9* and *GFP* were similarly inactive in mESC and induced during NPC differentiation (**Fig. 1C**). We then performed RCMC in NPCs from a clonal cell line with the re-introduced Sox9 full promoter and for cells carrying the Hyg-TK landing pad. Using the re-introduced Sox9-GFP transgene as virtual 4C viewpoint showed that the Sox9-FP transgene interacted exactly like the wildtype promoter with its entire TAD in mESCs and NPCs (**Fig. 1D, Fig. S1**). Notably, the same analysis with the Hyg-TK selection cassette showed that the interaction pattern with NPC-specific Sox9 enhancers resembled that of the Sox9 promoter (**Fig. 1D, Fig. S1C,D**). The interaction strength with the Sox9 TAD boundaries and intra-TAD CTCF sites was reduced, consistent with the lack of a CTCF site in the Hyg-TK selection cassette compared to the Sox9 promoter. These results validated that the Promoter-RMCE setup can be used to test the ability of promoters to respond to distal enhancers distributed throughout *Sox9* TAD.

We next sought to determine which specific regions within the *Sox9* promoter-proximal region might determine responsiveness. The 2.8kb *Sox9* full promoter contains three ENCODE-annotated proximal enhancer-like signature elements (pELS) as well as a constitutive CTCF binding site 2.3kb upstream of the TSS (**Fig. 1B**). We created a Promoter-RMCE library containing the *Sox9* full-length and core promoters as well as four truncated *Sox9* promoters. These lacked the CTCF site (ProxΔCTCF, 2.2kb) or additionally one or more of the annotated pELS elements: Prox-1.6, Prox-1.0, and Prox-0.5 (**Fig. 1D**). The CMV promoter is a 204bp element known to be frequently epigenetically silenced upon genomic integration^54–56^ in mESCs and derived cells. Therefore, we used this proximal promoter as a reference for an inactive promoter in all experiments. We integrated this truncated *Sox9* promoter library into the *Sox9* Promoter-RMCE mESC line, generating a pool of mESCs, each with a different promoter reporter integrated at on the modified *Sox9* allele. We differentiated this mESC pool into NPCs and quantified promoter activity by taking the ratio of normalized barcode abundance in cDNA vs gDNA using Amplicon-seq (**Fig. 1C**, Methods) ^57^.

The results of this genome-integrated reporter assay showed that the full length *Sox9* promoter drove the strongest expression while the core promoter alone showed no activity. The truncated *Sox9* promoters drove expression levels in between the full and core promoters (**Fig. 1E**). Removal of CTCF binding site caused 30.5% loss in activity, which did not decrease significantly after removal of pELS3 (Prox-1.6, 38.8% loss) or pELS2+3 (Prox-1.0, 33.3%). Additional deletion of pELS1 (Prox-0.5) resulted in a further reduction to 42.1% of the *Sox9* full promoter levels, rendering it non-significantly active compared to CMV. However, the activity of this Prox-0.5 promoter was still significantly higher than that of the inactive *Sox9* core promoter. These results show that *Sox9* full promoter responsiveness is conferred by different parts of the promoter-proximal region. While the CTCF site contributes ~30% to promoter activity, it is not the sole determinant of promoter-responsiveness. The region ~1kb to 66bp upstream of the TSS are required for the remaining ~70% of activity as measured in our assay.

### Full promoters show distinct autonomous activities in episomal and genome-integrated reporter assays

Since the Promoter-RMCE platform allows comparison of many promoters at the same genomic position, we next wanted to test whether any promoter would be activated by the *Sox9* regulatory domain. For this we compiled a library of 18 full promoters (**Fig. 2A**) from a range of genes with distinct expression patterns in mESCs and NPCs (**Fig. 2B**): housekeeping genes (*hPGK*, *Snx3*, *Arhgap1*); mESC-expressed genes (*TDGF1*, *Lefty1*, *Sox2*); NPC markers (*Sox9, Pax6, Vim, Nefm)*, developmental genes (*Tbx5, Gata4, Eomes, Hey2, Fgfr2),* as well as *Kcnj2* and *Kcnj16*, two genes located in the TAD upstream of the *Sox9* TAD^22,26^. The full promoter sequences reflect a broad range of chromatin accessibilities in NPCs (**Fig. 2C**), CpG densities (**Fig. 2D**), known core promoter motifs (**Fig. S3A**), as well as proximal promoter lengths (based on ENCODE-annotated pELS) (**Fig. 2E, Fig. S3A**). The promoters were cloned as a Promoter-RMCE compatible plasmid library comprising 90 elements in total.

**Figure 2:**
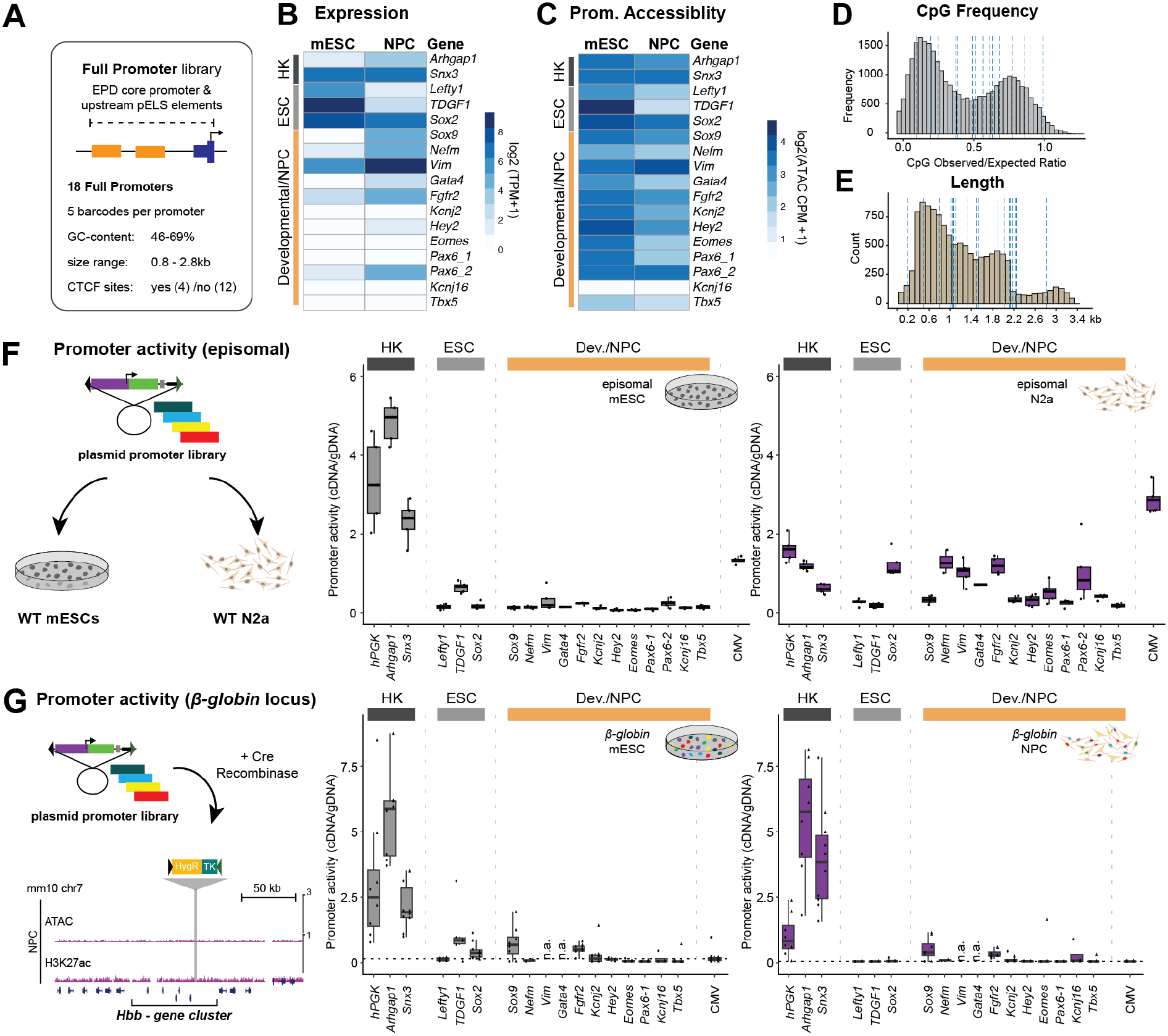
Full promoters show distinct autonomous activities in episomal and genome-integrated reporter assays. (**A**) Design of the sequences used for the construction of the full promoter library. (**B–E**) Characterization of the full promoter library showing expression. HK = housekeeping gene (**B**) and DNA accessibility (**C**) in their native loci in mESCs and NPCs, CpG frequency (**D**), and promoter length (**E**) Dashed blue lines = selected full promoters. (**F**) Activity of the full promoter library in episomal reporter assays in mESC and Neuro2a cells (**G**) NPC ATAC-and H3K27ac ChIP-seq tracks surrounding Promoter-RMCE integration site at the *β-globin* locus. Promoter-RMCE activities of the full promoters in mESC and NPCs. Triangles and dots = barcode values from two biological replicates. Dashed line: median CMV activity. n.a. = not available (not integrated in the genome).

To assess the intrinsic promoter activity, we first used this promoter library in an episomal reporter assay in mESCs and the neuroblastoma Neuro-2a (N2a) cell line, since mESC-derived NPCs did not yield sufficient transfection efficiencies (**Fig.2F**). In mESCs, housekeeping promoters and *CMV* showed strong activity, while all other promoters drove only weak or no expression in mESCs. By contrast, in N2a cells housekeeping and several other full promoters drove comparably strong expression. Others, including *Sox9*, drove weaker expression and only some showed no activity at all (**Fig. 2F**).

To test the autonomous activity of the full promoters as genome-integrated transgene, we generated a control Promoter-RMCE mESC line carrying the landing pad at the *β-globin* locus. This locus is active only in erythroid cells and part of a lamina-associated domain (LAD) in both mESCs and NPCs^58^. In our setup, this position served as a genomic control region devoid of distal enhancers in mESCs and NPCs (**Fig.2G**). After successful integration of the promoter library, we measured promoter activity in mESCs and derived NPCs. In contrast to the episomal assays, full promoter activity between mESCs and NPCs was very similar. Housekeeping promoters drove strong expression in both cell types, showing that the repressive LAD environment can be overcome by full promoter constructs. However, of the 15 remaining promoters only a few drove any expression in either mESCs or NPCs. The mESC promoters *TDGF1* and *Sox2* drove low expression in mESCs but were fully inactive in NPCs, while *Sox9* and *Fgfr2* promoters drove weak expression in mESCs and NPCs. All other promoters were inactive in both, mESCs and NPCs (**Fig. 2G**).

Together, these results show that full promoters contain cell-type specific autonomous regulatory elements whose activity can be measured in episomal reporter assays. The activity at the *β-globin* locus confirms the strong autonomous activity of housekeeping full promoters and shows that no other full promoters in our library encode a similar autonomous activity driving expression in mESCs and NPCs when integrated at that position.

### Promoter-RMCE reveals differential promoter responsiveness to the *Sox9* regulatory domain

We next used RMCE to integrate the full promoter library into the *Sox9* Promoter-RMCE mESC line and measured promoter activity in mESCs and NPCs (**Fig. 3A**). In mESCs, only few promoters drove strong expression from the *Sox9* position, namely the two housekeeping promoters *hPGK* and *Arhgap1* as well as the mESC promoter *TDGF1*, which was the most active. The housekeeping *Snx3* promoter and the mESC-specific *Lefty1* and *Sox2* promoters drove weak expression, as did the *Sox9* promoter. All other promoters, including CMV as a negative baseline, drove no expression in mESCs (**Fig. 3B**).

**Figure 3:**
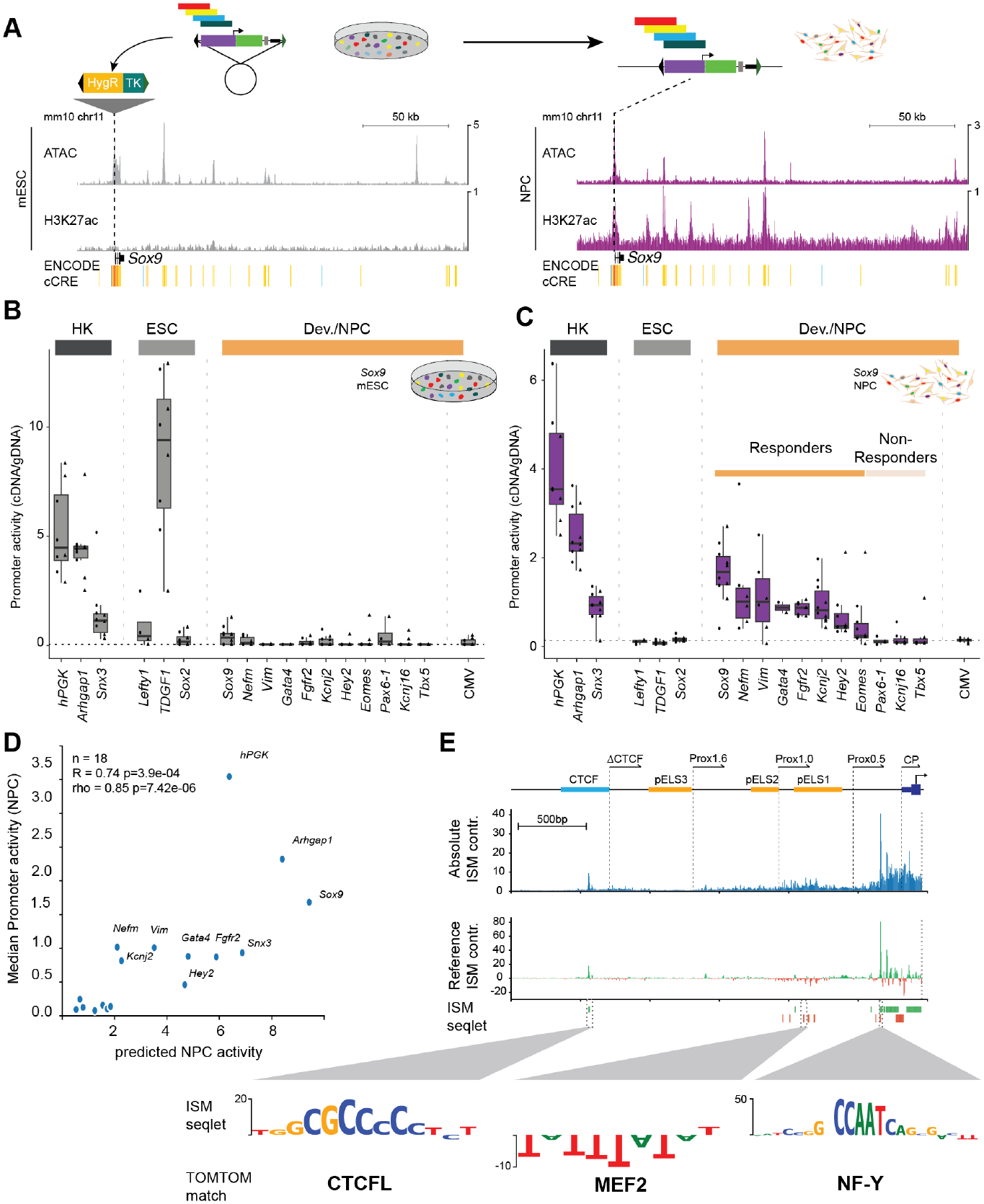
Promoter-RMCE shows differential full promoter activity at the *Sox9* locus in NPCs. (**A**) ATAC-and H3K27ac ChIP-seq at the *Sox9* locus in mESCs and NPCs. (**B,C**) Quantification of Promoter-RMCE activity at *Sox9* locus in mESC (**B**) and NPCs (**C**) Responders in C show activity above median CMV levels (dashed line) (**D**) Comparison of predicted promoter activity NPCs (fine-tuned Borzoi model, x-axis) to the measured promoter activity levels (y-axis). Pearson *r* and Spearman rho values are shown. (**E**) ISM-based contribution scores (CS) for the Sox9-FP. (top) absolute ISM-based CS, calculated by adding up the absolute nucleotide contribution values for all possible alleles at each position (middle) CS for the reference construct sequence. Green are positive, red are negative contribution values. (bottom) Called seqlets, colored by the positive/negative contribution. (inset) Logos of three example seqlets, nucleotide sizes correspond to CS and matched PWMs from the JASPAR database annotated below.

In contrast, full promoters in NPCs drove a wide range of expression levels that were decidedly distinct from their activity at the *β-globin* locus or in the episomal assay. First, housekeeping gene promoters drove stronger expression than most other promoters. Second, none of the three mESC-specific gene promoters drove any expression in NPCs. Third, the remaining 12 promoters produced a gradient of activity, from expression levels comparable to housekeeping promoters to non-detectable expression. Based on this separation we classified promoters in two groups: Responders and Non-responders (**Fig. 3C**). Within Responders, the *Sox9* promoter showed the highest activity, consistent with its endogenous role as part of this cis-regulatory domain. Two other NPC-specific promoters (*Nefm, Vim*) drove similarly high expression, followed by several other developmental gene promoters (*Gata4, Fgfr2, Kcnj2*, *Hey2*, *Eomes*) driving lower expression levels. In addition to the three mESC promoters, *Tbx5, Pax6-1, and Kcnj16* comprised the Non-responders, which all did not drive expression above CMV levels.

The differential activity of *Kcnj2* and *Kcnj16* mimicked the effect of large inversions at the *Sox9*locus, which causes *Sox9*-like misexpression *in vivo* of *Kcnj2* but not of the adjacent *Kcnj16*, despite both genes engaging with rearranged *Sox9* enhancers ^22,59^. Surprisingly, the promoter of the *bona fide* NPC marker *Pax6* included in our library was inactive in NPCs. We selected the *Pax6-1* promoter since it exhibits the highest signal in aggregated CAGE data. However, chromatin marks indicated that an alternative *Pax6-2* promoter located more proximal to the first coding exon might be the active promoter in in NPCs (**Fig. S4C**). We therefore included this alternative *Pax6-2* full promoter in a replicate experiment. Indeed, *Pax6-2* drove expression similar to other responding promoters, highlighting cell-type specific differences in alternative promoter responsiveness for the same protein-coding gene (**Fig. S4D**). Notably, several other promoters in our library also have alternative promoters. However, in contrast to *Pax6-2*, these do not show evidence of activity in NPCs, making them unlikely to act as Responders under tested conditions. In summary, these results show that full promoters display differential responsiveness to the *Sox9* cis-regulatory domain, with only some inducing transcription in response to distal enhancers during NPC differentiation, while others remain inactive.

### Chromatin state and general sequence features are weak predictors of promoter responsiveness

To test if any known molecular features might explain the observed promoter responsiveness, we correlated Promoter-RMCE activity at the *Sox9* locus with sequence and chromatin features of the native promoters in NPCs. The presence or absence of a CTCF binding site alone did not explain promoter responsiveness. While CTCF has been implicated in stabilizing enhancer– promoter interactions^14^, our dataset includes both responsive (*Sox9*, *Kcnj2*) and non-responsive (*Pax6-1*, *Lefty1*) promoters that contain constitutive CTCF sites, indicating that additional sequences within the promoter-proximal region are required to confer promoter responsiveness. Also, CpG density, endogenous gene expression level or endogenous promoter accessibility did not correlate with promoter responsiveness (**Fig. S3E-H**). Neither episomal activity nor full promoter activity at the *β-globin* and Sox9 locus correlated significantly with promoter activity at the Sox9 locus (**Fig. S3I,J)**. Only when removing the housekeeping promoters from the dataset, we found a significant correlation between the very weak activity at the *β-globin* locus and the strong activity at the Sox9 locus (**Fig. 3F, Fig. S3K**). However, the relative activity of the housekeeping promoters was distinct between both loci. Thus, while autonomous activity my play a part in promoter responsiveness, these results underline the importance of chromosomal position for the interpretation of a promoter’s regulatory function.

### A finetuned deep learning model predicts promoter responsiveness

To better understand the sequence determinants of promoter responsiveness, we finetuned a Borzoi machine learning model^60,61^ by low-rank adaptation (LoRA)^62^ and replacing the output head of the pretrained model with a new output head, trained using our mESC and NPC chromatin and RNA-seq data. Indeed, this finetuned model accurately predicted promoter responsiveness of the 18 full promoters integrated at the Sox9 promoter position (+/-250kb, Pearson’s R 0.74, Spearman Rho 0.85, (**Fig. 3D, Methods**)). To dissect which sequence elements drive this predictive power, we used in silico saturation mutagenesis^62^ across the full promoter sequences to calculate the bp contribution scores (CS). This revealed that rather than being confined to the core promoter, regions with high CS are distributed throughout the full promoter region (**Fig. 3E**). In many promoter-proximal regions sequences within ~1kb from the TSS harboured the majority of the informative signal (**Fig. S5**). However, additional high-contributing sequences were also present in more distal regions, highlighting the importance of different parts of the promoter-proximal region (**Fig. 3E, Fig. S5)**.

Intriguingly, the distribution of the CS was indicative of the truncated Sox9 promoter activity, correctly identifying the distal CTCF-motif and sequences between 800bp and 1.5kb as important for promoter activity (**Fig. 3E**). However, while the Borzoi-model accurately predicted the Sox9-CP to be inactive, it was unable to predict the stepwise loss of the truncated Sox9-promoters as measured by Promoter-RMCE (**Fig. S6A)**.

To understand which TFBS might drive the CS, we extracted contiguous regions of high CS (seqlets) using tangermeme^63^ and matched them to candidate TF motifs using Tomtom-lite^64^. Known promoter-associated motifs, most notably GC- and CCAAT-boxes, were deemed important in responding promoters. However, motif-scanning also identified e.g. GC-/CCAAT-boxes in non-responding promoters and simple enrichment analysis^65^ did not identify any top TF motifs in Responders (Supp. Data 1) to be enriched in the sequences of Responders vs. Non-responders. As indicated for the Sox9 full promoter (**Fig. 3E**), the CS for each full promoter instead appears to rely on a distinct combination of tissue specific TFBS and widely expressed activator motifs (GC-/CAAT-Boxes). Importantly, these high-contributing seqlets were not restricted close to the annotated core promoters but distributed across both core promoter and promoter-proximal region (**Fig. S5**), indicating that there is no canonical architecture underlying promoter responsiveness, rather each promoter appears to achieve its regulatory output through its own unique combinatorial sequence logic.

### Core promoters alone are insufficient to drive transcriptional response

Given this importance of chromosomal position for promoter activity, we next tested the ability of core promoters alone to drive expression at the *Sox9* position, especially since most MPRAs focus on the relatively short core promoter sequences individually or in combination with enhancers^9,10,12,28,29^. We cloned a library of 18 core promoters (based on ^66^; TSS +/-66bp) matching our set of full promoters and integrated these into mESCs. We then combined this mESC pool with the previously generated full promoter mESCs, to establish a new mESC pool of core and corresponding full promoters, and measured promoter activity in NPCs (**Fig. 4A**).

**Figure 4:**
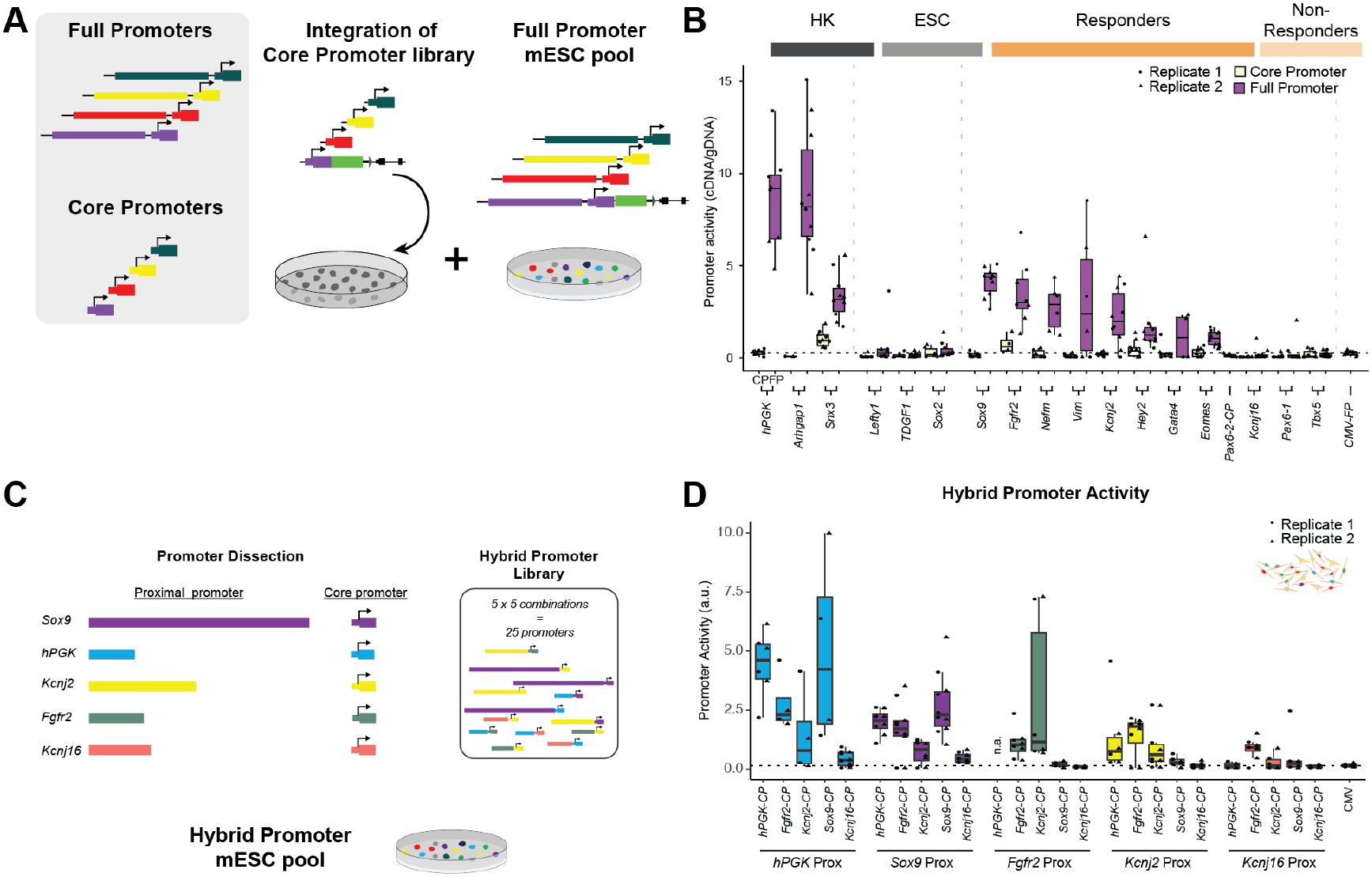
Core promoters are insufficient to respond to long-range activation but tune the responsiveness level of proximal promoter regions. (**A**) The core promoter library was integrated in a pool into mESCs and subsequently mixed with the full promoter mESC library prior to NPC differentiation. (**B**) Promoter-RMCE activity of corresponding core and full promoters in NPCs at the *Sox9* locus. Triangles and dots = barcode values from two biological replicates. Dashed line: median CMV activity. (**C**) Five proximal and core promoters were recombined in all iterations to create a hybrid promoter library. (**D**) Hybrid promoter activity in NPCs at the Sox9 locus grouped by proximal promoter identity. Triangles and dots = barcode values from two biological replicates. Dashed line: median CMV activity.

As expected, full promoters recapitulated our previous results. The core promoters, however, showed no or dramatically reduced activity in NPCs (**Fig. 4A**). Only two core promoters drove expression slightly above CMV levels, the housekeeping *Snx3* and *Fgfr2* (a Responder in the full promoter library). However, even the strongest core promoter expression was lower than that of the weakest responding full promoter (**Fig. 4A**). Together, this shows that core promoters by themselves are insufficient to drive gene expression, even at the *Sox9* genomic position that is tailored to receive signal from many distal enhancers. Thus, the promoter-proximal region must play the determining role in mediating promoter responsiveness.

### The core promoter modulates the transcriptional output of responsive proximal promoters

Finally, we hypothesized that while promoter-proximal regions determine the general responsiveness at a given locus, core promoters might have a role in tuning the transcriptional output. To test this experimentally, we designed and synthesized a library of 20 hybrid promoters, recombining core and proximal parts of 5 full promoters: *hPGK* (housekeeping), *Sox9*, *Kcnj2* and *Fgfr2* (Responders) and *Kcnj16* (Non-responder) (**Fig. 4B**). The NPC expression driven by these hybrid promoters together with their full promoter counterparts covered a wide range of activities consistent with our hypothesis, observations from previous experiments, as well as with predictions from the finetuned ML-model (**Fig. 4B, Fig. S6B**).

When grouped by proximal promoter, hybrid promoters containing the *hPGK* promoter-proximal region drove the highest expression, followed by *Sox9*, *Fgfr2*, *Kcnj2* and *Kcnj16* in descending activity (**Fig. S6G**). Within each group of proximal promoters, those combined with the *Kcnj16* core promoter drove the lowest expression levels and other core promoters relatively higher expression (**Fig. 4D**). The non-responding *Kcnj16* promoter-proximal region remained inactive when combined with all core promoters except for the *Fgfr2* core promoter. However, as the *Fgfr2* core promoter also drove expression when integrated by itself (**Fig. 4A**), the *Kcnj16:Fgfr2*-hybrid activity is likely due to the *Fgfr2* core promoter. In fact, when we grouped hybrid promoters by the identity of core promoter, *Fgfr2*-CP containing hybrid promoters showed the second-highest activity among developmental promoters included in the library (**Fig. S6H**). Importantly, those differential activites of core promoters are only observed in the presence of a proximal promoter, as core promoters alone fail to confer measurable enhancer responsiveness in our system. Thus, while core promoters differ in their intrinsic strength, this variation becomes functionally relevant only once responsiveness is established by the promoter-proximal region. Taken together, these findings support a model in which the promoter-proximal region determines whether a promoter can respond to distal enhancers, whereas the core promoter controls the transcriptional output.

## Discussion

Here we demonstrate promoter-specific responsiveness to long-range neuronal enhancers in the *Sox9* regulatory domain. We show that this specificity is determined by the promoter-proximal region and not by the core promoter. The core promoter, however, appears to modulate the output level of responsive promoters. Our work addresses the so far elusive phenomenon of enhancer-promoter compatibility in mammals. Evidence from individual loci suggested enhancer-promoter compatibility, yet systematic reporter screens of enhancers and core promoters produced inconclusive results^10–12^. Our results suggest that the promoter-proximal region is crucial for the response to an enhancer-rich regulatory domain, such as the *Sox9* TAD.

The promoter-proximal region directly upstream of the core promoter, sometimes referred to as the proximal enhancer^44^, has been mainly investigated for its ability to act as an autonomous activator. However, especially in development and during cell differentiation, genes are regulated by multiple enhancers located in large cis-regulatory domains that often contain several genes with independent expression patterns. By designing a genome-engineering strategy tailored to this regulatory scenario, our results reveal a less canonical regulatory function for promoter-proximal regions; as determining factor for enhancer-promoter compatibility. This suggests a mechanism similar to the recently described *facilitator* elements, which potentiate the activity of a distal enhancer when located in between the distal enhancer and the TSS^31,33,37,67^. Therefore, in the context of its cis-regulatory domain, we imagine that the promoter-proximal region serves as the ultimate facilitator element for the gene it controls. In this scenario, cell-type specific activity can be encoded in distal enhancers while the promoter-proximal region filters and transmits this activity to the core promoter. To fulfill this function, some low degree of activity might be necessary to be responsive to long-range enhancers whenever the gene needs to be expressed, but it is not sufficient to autonomously drive near-endogenous expression. A previous study^11^ compared the activity of different core promoters downstream of a short enhancer at distinct genomic positions, and the authors reported a general scaling of core promoter activity in response to the genomic loci. While the positive correlation between the weak activity at the *β*-Globin and strong activity at the *Sox9* loci may argue in a similar direction, the differential activity of the housekeeping promoters argue against such a simple scaling mechanism and for synergy between the promoter-proximal region and distal enhancers, which is supported by the conserved contacts to distal CREs between the *Sox9* promoter and the Hyg-TK selection cassette. Future work with tailored assays will need to test the biochemical nature of such synergistic effects.

Mechanistically, our 3D chromatin interaction pattern as well as our analysis of CTCF sites in the proximal promoters reveal a relevant, but limited contribution of 3D chromatin structure or alternative enhancer-promoter looping for promoter responsiveness. Our sequence analysis using the finetuned Borzoi ML-model shows that no simple TFBS logic explains promoter responsiveness in our data, but highlights the importance of sequence motifs across the core promoter and promoter-proximal region for responsiveness. Complementary work (see accompanying paper by Masoura et al.) also finds a boosting, facilitator-like function in the promoter-proximal region upstream of two core promoters in *Drosophila*, emphasizing the need to understand cis-regulatory elements in their direct and locus-wide genomic context.

Our data shows that the core promoter alone is not sufficient to respond to distal enhancers, since no core promoter was highly active when integrated by itself at the *Sox9* position. However, as shown by our library of hybrid promoters, core promoters have a function in tuning the output level once it is made accessible by the promoter-proximal region. In *Drosophila*, housekeeping and developmental core promoters show very distinct enhancer responsiveness (see also Masoura et al.), which was shown to be determined by the recruitment of distinct co-activators to different core promoters ^66,68^. The scale of our assay does not allow such granularity. However, similar compatibilities between the specific promoter-proximal regions and core promoters likely contribute to the differential activity we observe in the hybrid promoters.

The vast majority of previous studies on enhancer-promoter compatibility was based on episomal MPRAs which, by design, lack any locus-specific information and are limited in enhancer and promoter length. In contrast, our Promoter-RMCE assay is designed to investigate how locus-specific long-range signal of an enhancer-rich regulatory domain is integrated by relatively large full promoters. The importance of genomic context and differences in reporter assay design might have contributed to contradictory results as reported by Bergmann et al.^10^ and Martinez-Ara et al.^12^, using shorter (−244bp/+20bp) or longer (−375bp/+75bp) promoter fragments for their combinatorial MPRAs, respectively. Finally, considering core and proximal promoters not primarily as autonomous activators in reporter assays but as selective filters within their genomic context may enable the development of transgenes that resist silencing and can take advantage of the cis-regulatory domains at their integration sites.

Taken together, our work demonstrates the need to understand the function of cis-regulatory elements within their local and regional genomic context and place the promoter-proximal region as crucial selective filter with a facilitator-like role in its *cis*-regulatory domain, ultimately conferring enhancer-promoter compatibility.

### Limitations of the study

The size of the full promoters in this study limited the library size we were able to achieve. As a consequence, we are unable to experimentally address the sequence basis of promoter responsiveness using many variants of responsive and non-responsive promoters. Furthermore, promoter-proximal CREs with a facilitator like function might also be located downstream of the TSS^69^, which we are unable to test in our transgene design. Moreover, due to the complexity of the multi-enhancer *Sox9* regulatory domain, our setup is not suited to explore the compatibility rules between any specific distal enhancer and promoters. Instead, our results focus on comprehensively contrasting the facilitator-like role of the promoter-proximal region to that of the core promoter.

As the Promoter-RMCE setup causes a heterozygous loss of *Sox9*, there is a systematic difference to unedited NPCs. However, the lower *Sox9* levels are compatible with normal NPC differentiation, and the condition is identical for all inserted promoters. Finally, we use the *β*-Globin locus as enhancer-free control region that - despite being devoid of repressive chromatin marks - is located in a LAD in mESCs and NPCs, which provides an overall repressive environment. Extended to a greater number of alternative, more permissive loci, Promoter-RMCE or similar assays should provide additional insights on how full promoters interact and respond to their genomic surrounding.

## Acknowledgements

We would like to thank the MPI-MG Sequencing Core Unit for their help at various points throughout the project and Matt Kraushar for the Neuro2a cell line. We thank the Walk-Up Sequencing service of the Broad Institute of MIT and Harvard. We thank members of the Ibrahim laboratory for helpful discussions; and Tugce Aktas and Michael Robson for comments on the manuscript. D.M.I. was supported by funding from the DFG SPP 22.02 “3D Genome Architecture in Development and Disease” (IB139/1-1 and IB 139/6-1). Work in the Ibrahim Lab is supported by an ERC Starting Grant SYNREG (101076709). B.M., and S.M. were supported through the Marie Skłodowska-Curie Innovative Training Network (grant no. 813327 ‘ChromDesign’) under the European Union’s Horizon 2020 research and innovation program. Research in the laboratory of A.R.K. is supported by core funding from the EMBL, Deutsche Forschungsgemeinschaft (KR 5247/1-1, KR 5247/1-2, KR 5247/1-3), and the ERC (TFCoop-101125530). A.S.H. acknowledges funding support from the Novo Nordisk Foundation (NNF21SA0072102) and the G. Harold and Leila Y. Mathers Foundation.

## Author Contribution

B.M. designed and performed the experiments, analyzed the data and drafted the manuscript. D.M.I. conceived the study, designed the experiments, analyzed the data and wrote the manuscript. M.H. performed the RCMC experiments and analyzed the data together with A.S.H.. E.D.A. performed the machine learning and TFBS analyses. S.M. acquired funding, discussed results and contributed to the manuscript. A.K. supported the RMCE design, analysed the data and contributed to the manuscript. All authors read and approved the manuscript.

## Data Availability Statement

Sequencing data have been deposited to GEO under the GEO Accession number GSE336356 and can be accessed using the reviewer token abopaccebzgvngp. RCMC-data are submitted under GSE338328, reviewer token utkxoyuwzfgzlat.

## Methods

### Cell culture

Experiments were conducted in mouse G4 ESCs (XY, 129S6/SvEvTac x C57BL/6Ncr F1 hybrid) which were grown initially on mitomycin-inactivated CD1 mouse embryonic fibroblast feeder monolayer on gelatinised dishes at 37°C, 7.5% CO_2_. Wild type and mutant ESCs lines were cultured in ESC medium containing knockout DMEM with 4.5 mg/ml glucose and sodium pyruvate supplemented with 15% FCS, 10 mM Glutamine, 1x penicillin/streptomycin, 1x non-essential amino acids, 1x nucleosides, 0.1 mM beta-Mercaptoethanol and 1000 U/ml LIF. In days preceding nucleofection mutant cell lines were feeder depleted by plating on gelatin-coated dishes for 45 minutes after which they were transferred to a fresh gelatin coated dish and grown to full confluency.

### NPC differentiation

NPC differentiation was carried out as described^52^. In short, feeder depleted wild type or landing pad cell lines used for Promoter-RMCE were trypsynized, counted and washed with PBS. Each replicate of NPC differentiation was done using 4×10^6^ mESCs and cultured in 15 ml CA cell culture medium (DMEM supplemented with 10% FCS, 2 mM L-glutamine, 1 × non-essential amino acids and 5 μl per 500 ml β-mercaptoethanol) on bacteriological Greiner Petri dishes for 4 days with medium change after 2 days. After 4 and 6 days of cellular aggregation medium was changed to CA medium with addition of retinoic acid at final concentration of 5uM (#R2625, Sigma-Aldrich). Cellular aggregates were dissociated and collected after 8 days into several aliquots of 2 x 10^6^ cells for DNA and RNA extraction.

### Generation of landing pad cell lines at *Sox9* and *Hbb* loci

All steps leading to generation of landing pads mutant cell lines were performed using CRISPR as described previously^70^. To generate landing pad cell line at the *Sox9* locus, ESCs underwent sequential rounds of CRISPR: firstly CRISPR-mediated deletion of *Sox9* full promoter on one allele which was then followed by CRISPR-mediated knock-in. In short, ESCs were transfected with 4ug of two sgRNAs targeting *Sox9* full promoter using FuGENE HD (#E2311, Promega) according to manufacturer’s protocol. After 24 h, transfected cells were transferred on puromycin-resistant DR4 feeders and treated with puromycin for 48 h. Subsequently, ESCs grew for an additional 2-4 days after which clones were picked and transferred onto CD1 feeders in 96-well plates. 2-3 days later plates were split into three plates: 2 plates were frozen and 1 plate was further expanded and collected for DNA isolation followed by genotyping. After identification of clones that underwent deletion by breakpoint verification, positive clone was expanded and subjected to site-specific integration of hygromycin-thymidine kinase selection cassette. Here, ESCs were targeted with 8ug of sgRNA and 4 ug of the homology construct followed by the same procedure of selection and expansion as above. To generate landing pad cell line at the *β-globin* locus, ESCs were subjected exclusively to site-specific integration of selection cassette using 8µg of sgRNA and 4µg of the homology construct. A list of sgRNAs, primers used to amplify homology arms from the mouse genome and primers used to genotype resulting mESC lines can be found in Supplementary Table 1.

### WT full promoter library design and cloning

To determine sequences of full promoters we retrieved experimentally validated TSSs deposited in EPD Eukaryotic Promoter Database (*Mus musculus*, mm10^71^). We defined the region of the full promoter sequence by incorporating information provided by ENCODE Project Consortium^44^. With the exception of the *Sox9* promoter, we designed full promoter to span from ~66bp downstream of TSS and encompass all proximal enhancer-like signatures (high DNase and H3K27ac with low H3K4me3 signal within 2kb of TSS). In the case of the *Sox9* full promoter, the sequence was extended by ~1kb upstream of pELS3 to incorporate the CTCF binding site. Each promoter sequence was PCR amplified from the mouse genomic DNA (with exception of hPGK promoter coming from human genome which was amplified from available reporter vector) using S7 Polymerase (#MD-S7-500, Mobidiag) and using following PCR conditions: 98°C for 30 s; 33 cycles of 98°C for 10 s, 60 - 64°C for 30 s (depending on the sequence) and 72°C for 1-2 mins (depending on the sequence length). PCR primers were designed for each promoter separately, providing overhangs suitable for Gibson cloning into receiving donor plasmid, PacI and SphI-HF restriction sites and 15 random base pairs making the barcode sequence. PCR products were gel purified and cloned into ClaI & SalI digested BMV1.1 donor plasmid backbone using HiFi DNA Assembly Master Mix (#E2621L, NEB) following manufacturer’s protocol. Gibson reactions were purified and transformed by electroporation into MegaX DH10B T1^R^ Electrocomp™ Cells (#C640003, Invitrogen) which are suitable for library construction applications. Subsequently, for each promoter at least 6 colonies were picked, plasmid DNA was isolated and sent for Sanger sequencing to check sequence integrity and determine barcode sequence. Each cloned promoter was associated with 5 unique barcodes to provide measurement robustness and to control for barcode-specific effects. Subsequently, correct promoter constructs were digested with PacI and SphI-HF, gel purified and an eGFP sequence was cloned between promoter and the barcode using T4 DNA Ligase following manufacturer’s manual (#M0202, NEB). All promoter – eGFP construct plasmids were pooled equimolarly to generate plasmid donor library, retransformed again into MegaX DH10B T1^R^ Electrocomp™ Cells and grown overnight in liquid culture. Plasmid DNA was extracted on the following day using NucleoBond Xtra MIDI EF (#740420.50, Macherey-Nagel) and used for RMCE transfection into mESCs.

Promoter genomic coordinates, primers used to amplify promoters and a list of barcodes associated to promoters are provided in Supplementary Table 1.

### Core promoter cloning

The core promoter library encompasses core promoters (−66 to +66 bp around EPD-annotated TSS) corresponding to promoters in full promoter library. We ordered the library as a single-stranded oligo pool from Twist Bioscience, where each promoter is present in the library 5 times, each time with a different pre-determined barcode. Similar to the full promoter library, each primer sequence was designed to have common primer-binding sites being the overhangs for Gibson cloning into donor vector and restriction sites for eGFP cloning. Firstly, to avoid overamplification, the oligo pool was amplified by setting up 8 PCR reactions with KAPA HiFi HotStart PCR Kit (#2502, KAPA) using 10 ng of DNA and following the protocol: 95°C for 3 min; 10 cycles of 98°C for 20 s, 60°C for 15 s and 72°C for 15 s and final extension at 72°C for 1 min. Then, PCR reactions were pooled, purified with AMPure beads and cloned into ClaI & SalI digested BMV1.1 plasmid backbone using HiFi DNA Assembly Master Mix (#E2621L, NEB) following manufacturer’s protocol. Gibson reactions were purified and transformed by electroporation into MegaX DH10B T1^R^ Electrocomp™ Cells. Promoter constructs were then digested with PacI and SphI-HF, gel purified and eGFP sequence was cloned between promoter and the barcode using T4 DNA Ligase following manufacturer’s manual. The pool of construct-eGFP plasmids was then transformed into bacteria, incubated with shaking overnight at 37°C and plasmid DNA was extracted on the following day using NucleoBond Xtra MIDI EF (#740420.50, Macherey-Nagel).

### Hybrid promoter library design and cloning

Hybrid promoter constructs were ordered as clonal genes from Twist Bioscience with the exception of *Kcnj2:Fgfr2* and *Sox9:Fgfr2* which were ordered from Azenta GeneWiz. The sequences of the constructs were designed based on core promoter (−66 to +66 bp around EPD-annotated TSS) and full promoter sequence definition with small sequence modifications allowing for efficient synthesis. Synthesized constructs were amplified using KAPA polymerase using following PCR conditions: 95°C for 3 min; 33 cycles of 98°C for 20 s, 72°C for 15 s and 72°C for 3 mins. PCR reactions were purified using NucleoSpin Gel and PCR Clean-up Kit (#11992242, Macherey-Nagel). Purified hybrid promoter constructs were cloned into ClaI & SalI digested BMV1.1 plasmid backbone using HiFi DNA Assembly Master Mix (#E2621L, NEB) following manufacturer’s protocol. Gibson reactions were purified and transformed by electroporation into MegaX DH10B T1^R^ Electrocomp™ Cells. Subsequently, for each hybrid promoter construct at least 6 colonies were picked and plasmid DNA was isolated using NucleoSpin Plasmid Mini Kit ($740588.50, Macherey-Nagel) and sent for Sanger sequencing to check sequence integrity and determine barcode sequence, aiming at getting 5 different barcodes per construct. After that, correct promoter constructs were digested with PacI and SphI-HF, gel purified and eGFP sequence was cloned using T4 DNA Ligase following the manufacturer’s manual (#M0202, NEB). Hybrid promoter – eGFP construct plasmids were pooled equimolarly to generate plasmid donor library used for RMCE transfection into mESCs. All hybrid promoter sequences and primers used to amplify them (reused from cloning WT full promoter library) are available in Supplementary Table 1.

### Variant *Sox9* promoter library design and cloning

Lengths of variants of *Sox9* promoters were determined based on ENCODE annotations. Variants were amplified from the mouse genome using primers designed to bind to specific genomic sequence and provide overhangs suitable for Gibson cloning into receiving donor plasmid, PacI and SphI-HF restriction sites and 15 random base pairs making the barcode sequence. PCR was performed using S7 Fusion S7 Polymerase (#MD-S7-500) and using following PCR conditions: 98°C for 30 s; 33 cycles of 98°C for 10 s, 64°C for 30 s and 72°C for 1-2 mins (depending on the variant length). Following the PCR, purification, cloning into receiving vector, determining barcode sequence, cloning the eGFP and transforming into bacteria was done as described above for WT FP library or hybrid promoter library. All primers used to amplify variant Sox9 promoters from the genome are available in Supplementary Table 1.

### Integration of promoters into landing pad cell lines and selection of positive clones

The cell lines containing the Hygromycin-Thymidine Kinase selection cassette were cultured in ESC medium described above for 2-4 days followed by 10 days of selection with hygromycin (140µg/ml) to ensure expression of the Hyg-TK transgene for Ganciclovir selection. Cells were then feeder depleted and expanded in the days preceding transfection aiming at getting at around 24 to 50×10^6^ cells, depending on experiment. On a day of transfection, cells were collected by trypsinization and counted and divided accordingly into 15mL Falcon tubes. For WT Full Promoter library, Core Promoter library and Hybrid Promoter library 2 biological replicates were performed, each replicate required 24×10^6^ cells divided into 6 transfection reactions. Due to lower number of constructs integrated for variant *Sox9* Promoter library, each of two biological replicates was done using 12×10^6^ cells divided into 3 transfection reactions. In short, in each transfection reaction 4×10^6^ cells were nucleofected with 25ug of promoter plasmid library and 15ug of plasmid Cre recombinase using P3 Primary Cell 4D Nucleofector X Kit L following manufacturer’s protocol (#13429329, Lonza). Immediately after, cells were transferred to 15 cm cell culture dish with 25mL of ESC medium supplemented with LIF. After two days of post-transfection recovery (with ESC medium + LIF changed daily), cells were cultured in ESC medium +LIF, additionally supplemented with ganciclovir to select for cells that underwent recombination. Here, for cells with the LP at the *Sox9* locus ganciclovir at 10uM concentration was used, while the cells with LP at the *β-globin* were selected with 3uM ganciclovir. The selection lasted 14 or 8 days, respectively. After the selection process, cells were expanded and collected either into several aliquots of 2 x 10^6^ cells for DNA and RNA extraction or to start NPC differentiation process.

### Episomal reporter assay in mESCs and N2a cell lines

For the episomal reporter assay, mESCs and N2a cells were transfected using P3 Primary Cell 4D Nucleofector XL Kit (Lonza, #13429329). For each cell type, 3 biological replicates were performed by transfecting 1 ×10^6^ cells with 3 ug of full promoter library per replicate. After 24 h, RNA was harvested using the RNeasy Mini Kit Column (Qiagen, #74104), including on-column DNase treatment for 15 min. Both input plasmid library and isolated RNA were in the next step subjected to preparation of amplicon sequencing libraries.

### Preparation of amplicon-seq libraries

For gDNA amplicon-seq, DNA from collected mESC or NPC RMCE libraries was isolated using Puregene Cell Kit (#158043, QIAGEN). Each biological replicate (1x for mESC samples and 2x for NPC samples) was prepared in two technical replicates. For each technical replicate 8 PCR reactions (each 200ng gDNA) were set up to amplify barcode sequences using S7 Fusion Polymerase kit (#MD-S7-500, Mobidiag) with following PCR conditions: 98°C for 30 s; 30 cycles of 98°C for 10 s, 68°C for 30 s and 72°C for 30 s and using NEBNext® Multiplex Oligos for Illumina®, Index Primer Set 1-4 (NEB). For any given technical replicate, all 8 PCR reactions were pooled and purified together using AMPure beads.

For cDNA amplicon-seq, RNA from collected mESC or NPC RMCE libraries was isolated using Rneasy Mini Kit (#74104, QIAGEN) with on-column gDNA digestion following the manufacturer’s protocol. For each biological replicate 4ug of RNA was reverse transcribed into cDNA using PrimeScript™ RT Reagent Kit with gDNA Eraser (#RR047A, Takara). Next, PCR reactions were set up using 200ng of cDNA per reaction, pooled and purified as described above for gDNA amplicon-seq.

All eluted libraries were checked using High Sensitivity D1000 ScreenTape Assay ((#5067-5584 and #5067-5585, Agilent) and sequenced on NovaSeq2, NextSeq2000 (Illumina) or AVITI (Element Biosciences).

### ATAC-seq library preparation

ATAC-seq was performed as described previously^72^. Briefly, 6×10^5^ mESCs or NPCs were used for each replicate. First, cells were washed with cold PBS. After centrifugation, cells were lysed in fresh lysis buffer (10mM TrisCl pH7.5, 10mM NaCl, 3mM MgCl2, 0.1% (v/v) Igepal CA-630) on ice for 3 minutes. Tn5 transposition was carried out for 30 minutes at 37°C and DNA was later purified with MinElute Reaction Cleanup kit (#28204, Qiagen). Nextera indexing primers were added during library amplification from purified DNA using NEBNext High-Fidelity 2X PCR Master Mix (#M0541S, NEB). The number of cycles used to amplify library was determined by qPCR. After double-sided size selection, size distribution of the libraries was determined with High Sensitivity D1000 ScreenTape Assay (#5067-5584 and #5067-5585, Agilent) aiming for nucleosomal pattern and then ready libraries were sequenced on NovaSeq2 (Illumina) with 100bp paired-end reads. ATAC-seq experiments were performed in duplicates.

### ChIPmentation

ChiPmentation libraries were prepared using iDeal ChIP-seq kit for Histones (#C01010051, Diagenode) followed by Tn5 tagmentation and library amplification. Briefly, NPCs were dissociated and fixed with 1% MeOH-free formaldehyde (#28906, Thermo Scientific) in PBS on ice for 10 minutes. Fixed cells were quenched using glycine and snap frozen until proceeding with further steps. Next, cells were resuspended in lysis buffer iL1b and incubated for 20 minutes at 4°C on a rotating machine. After pelleting cells at 500 rcf for 5 minutes, they were resuspended in lysis buffer iL2 and incubated again at 4°C for 10 minutes. After centrifugation (500 rcf, 5 minutes) cells were resuspended in Sonication buffer and the chromatin was sheared using standard High-Power Bioruptor settings: 30 cycles (30 seconds “ON”, 30 seconds “OFF”. Sonication was interrupted every 5 cycles to centrifuge the liquid in the tubes. After final cycles sheared chromatin was transferred to new 1.5mL tubes and centrifuged at 16000 rcf, 4°C for 10 minutes. ProtA beads were blocked with appropriate antibodies and after 4 hrs at 4°C 1-2ug of sheared chromatin was added to each sample. After overnight incubation with antibodies, samples were washed with buffers iW1-iW4. Tagmentation of pulled-down chromatin with Tn5 was carried out for 5 minutes at 37°C. Reaction was de-crosslinked with Proteinase K at 65°C overnight. DNA was then purified using AMPure beads at ratio 1.8X :1. Nextera indexing primers were added during library amplification from purified DNA with NEBNext High-Fidelity 2X PCR Master Mix (# M0541S, NEB) where the number of cycles to be used for final amplification was based on qPCR Ct values. After double-sided size selection, correct distribution of the library size was determined using High Sensitivity D1000 ScreenTape Assay (Agilent) and sequenced paired-end on NovaSeq2 (Illumina) or AVITI (Element Biosciences).

### RNA-seq library preparation

mESCs were trypsynized and feeder depleted for 45 minutes. WT and mutant NPCs were collected by trypsinization as described previously^52^. All collected samples were snap frozen and stored at −80°C. Total RNA was extracted using RNeasy Mini Kit following manufacturer’s protocol. 500ng of RNA was used for each sample to construct sequencing library using Kapa HyperPrep Kit. Samples were sequenced on a NovaSeq2 with 100bp paired-end reads. RNA-seq experiments were performed in duplicates.

### qRT-PCR

Following cell collection and RNA isolation with RNeasy Mini Kit, cDNA synthesis of 500ng of RNA was performed with SuperScriptIV RT (#18090010, ThermoFisher Scientific). Relative quantification was performed using Design and Analysis Software by normalization to *GAPDH* and *Rps9* expression. qPCR primers used in the study are listed in Supplementary Table 1.

### Region Capture Micro-C library preparation

Region Capture Micro-C (RCMC) was performed as described previously^17^. 3×10^7^ mESCs or NPCs were used for the Sox9-FP transgene clonal cell line, while 6×10^7^ NPCs were used for cells containing the landing pad sequence which had undergone RMCE transfection. Cells were dissociated, washed with PBS and aliquoted into lots of 3×10^7^ cells for crosslinking. Cells were crosslinked with gentle rotation for 35 minutes with 3 mM disuccinimidyl glutarate at room temperature. After 35 minutes of crosslinking, formaldehyde was added to a final concentration of 1%, and crosslinking was continued for a further 10 minutes. Crosslinking was quenched by addition of Tris pH 7.5 to 0.375 M and incubation for 5 minutes at room temperature. Crosslinked cells were pelleted, washed twice with PBS, and then aliquoted into tubes of 5×10^6^ cells each and snap frozen.

To determine the optimal MNase quantity for in situ chromatin digestion, an MNase titration was performed on a single 5×10^6^-cell aliquot of each condition. The aliquots were thawed on ice and incubated in buffer MB1 (10 mM Tris-HCl pH 7.5, 50 mM NaCl, 5 mM MgCl_2_, 1 mM CaCl_2_, 100 µg/mL BSA (Sigma-Aldrich, B8667), 1x Protease Inhibitor Cocktail (cOmplete, EDTA-free Protease Inhibitor Cocktail, Sigma-Aldrich, 11873580001), 0.2% NP-40 Alternative (Millipore Sigma #492018)) for 20 minutes. Cells were washed once more with MB1 and then split into 1×10^6^-cell aliquots for MNase (Worthington Biochemical, LS004798) digestion. Aliquots from the same sample were digested by addition of a range of MNase quantities between 2 and 20 U, and incubation at 37 °C for 20 minutes with constant shaking. MNase was quenched by addition of EGTA to 4 mM and heating at 65 °C for 10 minutes. Digested nuclei were pelleted and the supernatant removed before resuspending in reverse crosslinking mix (0.8X TE, 1 mg/mL Proteinase K (Viagen Biotech, 501-PK), 1% SDS, 0.1 mg/mL RNase A (ThermoFisher, EN0531), 200 mM NaCl) and incubating at 65 °C overnight constant shaking to reverse crosslinks and dissolve cell debris. DNA was purified using the Zymo DNA Clean and Concentrator (DCC-25) kit (Zymo Research, D4034). Purified DNA was run on a gel, and the optimal MNase quantity determined based on the relative intensities of mono-, di- and trinucleosomal bands. To prepare Micro-C libraries, samples, in aliquots of 5×10^6^ cells, were treated as in MNase titration up to the MNase digestion step. Samples were digested in 5×10^6^-cell aliquots, resuspended in 5x the volume as for 1×10^6^-cell aliquots during the titration, and with 5x the quantity of MNase determined from the titration. MNase digestion was quenched by addition of EGTA to 4 mM and heating at 65 °C for 10 minutes. Nuclei were pelleted and washed twice with MB2 (10 mM Tris-HCl pH 7.5, 50 mM NaCl, 10 mM MgCl_2_, 100 µg/mL BSA). Nuclei were then resuspended in 3’ end exonuclease reaction mix (1x NEBuffer 2.1 (New England Biolabs, B7202S), 2 mM ATP (Thermo Fisher, R1441), 5 mM DTT (Sigma-Aldrich, 10197777001), 0.5 U/µL T4 PNK (New England Biolabs, M0201) and incubated at 37 °C with constant mixing. After 15 minutes, Klenow fragment (New England Biolabs, M0210L) was added to 0.5 U/µL, and the samples were incubated with mixing at 37 °C for a further 15 minutes. Biotin labeling reagents (200 µM biotin-14-dATP (Jena Bioscience, NU-835-BIO14), 200 µM biotin-11-dCTP (Jena Bioscience, NU-809-BIOX), 200 µM dTTP (Jena Bioscience, NU-1004), 200 µM dGTP (Jena Bioscience, NU-1003), 1x T4 DNA ligase buffer (New England Biolabs, B0202S), 100 µg/mL BSA) were then added to the samples on ice. End labeling of digested fragments was performed by incubation at 25 °C for 45 minutes, with one minute on, three minutes off interval mixing at 1,000 rpm. The labeling reaction was quenched by addition of EDTA to 30 mM and incubation with interval mixing as before at 65 °C for 20 minutes. Nuclei were washed with MB3 (50 mM Tris-HCl pH 7.5, 10 mM MgCl_2_, 100 µg/mL BSA). After removal of MB3, nuclei were resuspended in proximity ligation mix (1x T4 DNA ligase buffer, 100 µg/mL BSA, 20,000 U/mL T4 DNA ligase (New England Biolabs, M0202S)) and proximity ligation was carried out by incubation overnight at RT with gentle mixing. Nuclei were pelleted and resuspended in a mix for removal of unligated DNA ends (1x NEBuffer 1 (New England Biolabs, B7001S), 5 U/µL Exonuclease III (New England Biolabs, M0206)) with interval mixing for 15 minutes at 37 °C. Samples were reverse crosslinked by addition of 2 mg/mL Proteinase K, 1% SDS, 0.1 mg/mL RNase A, and 200 mM NaCl and incubation overnight at 65 °C with 1,000 rpm shaking. DNA was purified using the Zymo DCC-25 kit, and then run on a 1% agarose gel. Dinucleosome-sized DNA fragments were excised from the gel and purified using the Zymoclean Gel DNA recovery kit (Zymo Research, D4008). Size-selected and gel-purified biotinylated DNA was pulled down using MyOne Streptavidin T1 Dynabeads (Invitrogen, 65601) suspended in bead wash (BW) buffer (5 mM Tris-HCl pH 7.5, 1 M NaCl, 0.5 mM EDTA) at RT overnight. Beads were washed twice with TWEEN BW (TBW) buffer (0.1% TWEEN 20, 5 mM Tris-HCl pH 7.5, 1 M NaCl, 0.5 mM EDTA) and once with 10 mM Tris-HCl pH 7.5 before resuspending in 0.1x TE buffer. Micro-C libraries were prepared from the Dynabead-bound DNA using the NEBNext Ultra II DNA Library Prep kit (New England Biolabs, E7645) according to the manufacturer’s protocol, with the addition of interval mixing during all incubations up to the end of adaptor ligation. After adaptor ligation and prior to PCR, beads were washed twice with TBW and once with 10 mM Tris-HCl pH 7.5. The number of cycles for PCR amplification were determined by running a test PCR on an aliquot of the main sample. Following PCR, Micro-C libraries were purified using 0.9x AmPure XP beads (Beckman Coulter, A63880) according to the library prep kit protocol.

To capture reads originating from the region around *Sox9*, a custom probe panel (Twist Bioscience, 101001) was designed with 80-mer probes covering the region between 109039512 and 114907512 on chromosome 11 (mm39). The custom probe panel also included probes matching the sequences of the full length promoter library and landing pad sequence to allow capture of reads from edited alleles. This custom panel was used with the Twist Target Enrichment Standard Hybridization v2 Protocol to generate RCMC libraries. The number of PCR cycles to use for final RCMC library amplification was determined by running a test PCR on an aliquot of the main sample. RCMC library concentrations were determined by qPCR and sequenced (100 cycles, paired-end) on the NovaSeq X sequencing platform using at the Broad Institute of MIT and Harvard.

### Quantification and analyses ATAC-seq analysis

Nextera Tn5 adaptor sequences were trimmed from fastq reads using *cutadapt* before further processing. Reads were aligned to mm10 genome using bowtie v2.3.5.1. Duplicated reads were removed using *MarkDuplcates* (Picard v2.23.4). Reads were sorted and filtered using samtools v1.10 to remove unmapped reads, low quality reads and mitochondrial reads. Filtered bam files from replicates were merged to calculate CPM for promoter regions (TSS −3/+2kb) using *bamCount* (bamsignals). ATAC-seq experiments were performed in duplicates. For differential accessibility analysis, peaks were called on merged BAM files using MACS2 (callpeak, -f BAMPE, -q 0.01) to generate a consensus peak set. Read counts over consensus peaks were analyzed using edgeR, with TMM normalization and a generalized linear model with likelihood ratio testing. Peaks with a false discovery rate (FDR) < 0.05 and an absolute log2 fold change > 1 were considered significantly differentially accessible.

### Amplicon-seq analysis

For each technical replicate of each library barcodes were extracted from the single end Illumina reads by using a custom script that extracts the 15 bp barcode sequence at the constant position of the read (28 – 42bp). All extracted barcodes in gDNA and cDNA were then associated to their corresponding promoters (Supplementary Table 1) and counted. Barcode counts from both technical replicates for each biological replicate were combined and only barcodes that had > 1 read in mESC and in both biological NPC replicates were retained for downstream analysis. Barcode counts were then normalized to the total number of barcodes reads from each sample. Promoter activity for any given barcode was calculated as cDNA/gDNA ratio (for promoter-RMCE) or as cDNA/plasmid DNA ratio (for episomal MPRA) of normalized barcode counts. To minimize the influence of extreme values on correlation analyses, observations within biological replicates activity above the 95th percentile were flagged as potential outliers. The most extreme of these were removed prior to plotting. Statistical comparisons between promoters and the low-activity CMV control were performed using two-sided Wilcoxon rank-sum tests at the barcode level. P-values were adjusted for multiple testing using the Benjamini– Hochberg procedure to control the false discovery rate. Promoters were classified as responders only if they showed both a significant difference from CMV (BH-adjusted p < 0.05) and a median fold-change greater than 1.5 relative to CMV.

### ChIPmentation analysis

ChIPmentation sequencing reads were aligned to the mouse reference genome (mm10) using Bowtie2^73^ with the --very-sensitive setting, a maximum insert size of 1000 bp, and duplicate marking by samblaster. Low-quality alignments (MAPQ < 10) and mitochondrial reads were removed, and duplicate reads were filtered out. BAM files were sorted and indexed with SAMtools^74^, and biological replicates were merged. For visualization, coverage tracks were generated in bigWig format using bamCoverage (deepTools2^75^) with CPM normalization.

### RNA-seq analysis

Paired-end 100 bp RNA-seq reads were mapped to the mouse reference genome (mm10) using STAR^76^ with splice junctions guided by GENCODE annotation. Mapping was performed with the following parameters: --outFilterMultimapNmax 20, --alignIntronMax 1000000. Resulting SAM files were converted to BAM, merged across biological replicates, sorted, and indexed using SAMtools. For visualization, normalized coverage tracks were generated in bigWig format using bamCoverage with RPKM normalization, bin size of 50 bp, and a minimum mapping quality of 30.

### Machine learning analysis

The pretrained Borzoi model^60^ was downloaded from gs://scbasset_tutorial_data/baskerville_transfer_mouse/pretrain_trunks/trunk_r1.h5. Finetuning was performed on the mm10 reference genome, using the same validation and test folds as in Borzoi. LoRA^62^ adapters with rank 8 were used for finetuning, using default parameters^61^. To obtain model predictions of our constructs within the genomic context, 524 kb sequences were created by extending construct sequences with the reference genomic sequences on the left and right. Predicted promoter activities were defined as the mean coverage predictions overlapping with the GFP position in the NPC RNA-seq track. Nucleotide contribution scores (CS) were computed using in silico saturation mutagenesis^77^, and model predictions were obtained for each possible mutation in the full promoter, using the same NPC GFP RNA expression readout as described above. Final CS for each allele were calculated by subtracting the mean expression across the four alleles. Seqlets were identified using recursive_seqlets from the tangermeme package^63^ (version 1.0.4), using a p-value threshold of 0.05, and additional_flanks=1. Seqlets were annotated with known TF binding motifs from the JASPAR 2026 core vertebrates non-redundant motifs^78^, using the Tomtom-lite^64^ implementation from memesuite-lite^79^ (version 0.4.0), using 0.05 p-value and 1.0 e-value thresholds. Motifs with at least 1 TOMTOM hit (e-val 0.2) for at least 5/11 Responders were selected for motif enrichment analysis. Then SEA^65^ enrichment with a p-value threshold of 0.05 was performed for these motifs.

### Region-Capture Micro-C Analysis

Paired-end sequencing reads from RCMC libraries were processed from alignment to mcool files using a previously developed analysis pipeline (https://github.com/ahansenlab/MicroC_RCMC_analysis). Data from clonal *Sox9*-FP cells were aligned twice: to the WT mm39 genome assembly, and to a modified genome in which the *Sox9* gene was replaced with the Sox9-FP transgene. To confirm the similarity of interactions from the *Sox9* locus, data were reamapped to a modified genome including a proxy ‘chromosome’ containing either the Sox9-FP transgene (WT genome) or the WT *Sox9* sequence (Sox9-FP transgene-modified genome). This caused reads arising from within the *Sox9* promoter, which is present on both alleles, to map at multiple sites and be discarded. To align data from cells containing the landing pad sequence which had undergone RMCE transfection with the full-length promoter library, a modified genome was created in which all endogenous copies of the full-length promoters included were deleted and reinserted as individual ‘chromosome’ sequences, and the WT *Sox9* gene was replaced with the Hyg-TK cassette landing pad. RCMC contact maps and other genomic tracks were visualized using a combination of coolbox^80^ (v0.3.8) and pygenometracks^81,82^ (v3.8)

Virtual 4C plots were generated from RCMC data using cooltools^83^ (v0.7.1) virtual4c at 1 kb resolution using the WT *Sox9* gene, Sox9-FP transgene, or Hyg-TK cassette as viewpoints. Mapping and normalization artefacts within plotted regions were masked using a combination of the ENCODE mm10 blacklist^84^ converted to mm39 using UCSC LiftOver^85^ (https://genome.ucsc.edu/cgi-bin/hgLiftOver) and a manually curated list of loci covering regions of low mappability, determined using cooltools coverage at 1 kb resolution.

2D pileups of contacts between the *Sox9* locus at ATAC-seq peaks within the region around *Sox9* gained in NPCs were generated from RCMC contact matrices using the coolpuppy^86^ API at 500 bp resolution. 1D pileups of the same contacts were generated from bigwigs of 1 kb-resolution virtual 4C data using computeMatrix reference-point and plotProfile from the deepTools^87^ suite. 1D pileups were generated with a window of +/− 10 kb up- and downstream of the peaks. Differential peaks overlapping or within 10 kb of *Sox9* (peaks 6 and 7) were excluded from pileup analysis. Due to the lower ability to resolve neighboring interaction peaks in 1D pileups, 1D pileups were also generated excluding differential peak 15 to avoid biasing the plot, as a result of its proximity to a CTCF site which appears stronger in NPC than mESC and in alleles with the CTCF-containing *Sox9* promoter.

