## Supplemental Figures for "Promoter-responsiveness to distal regulatory elements is mediated by the promoter-proximal region"

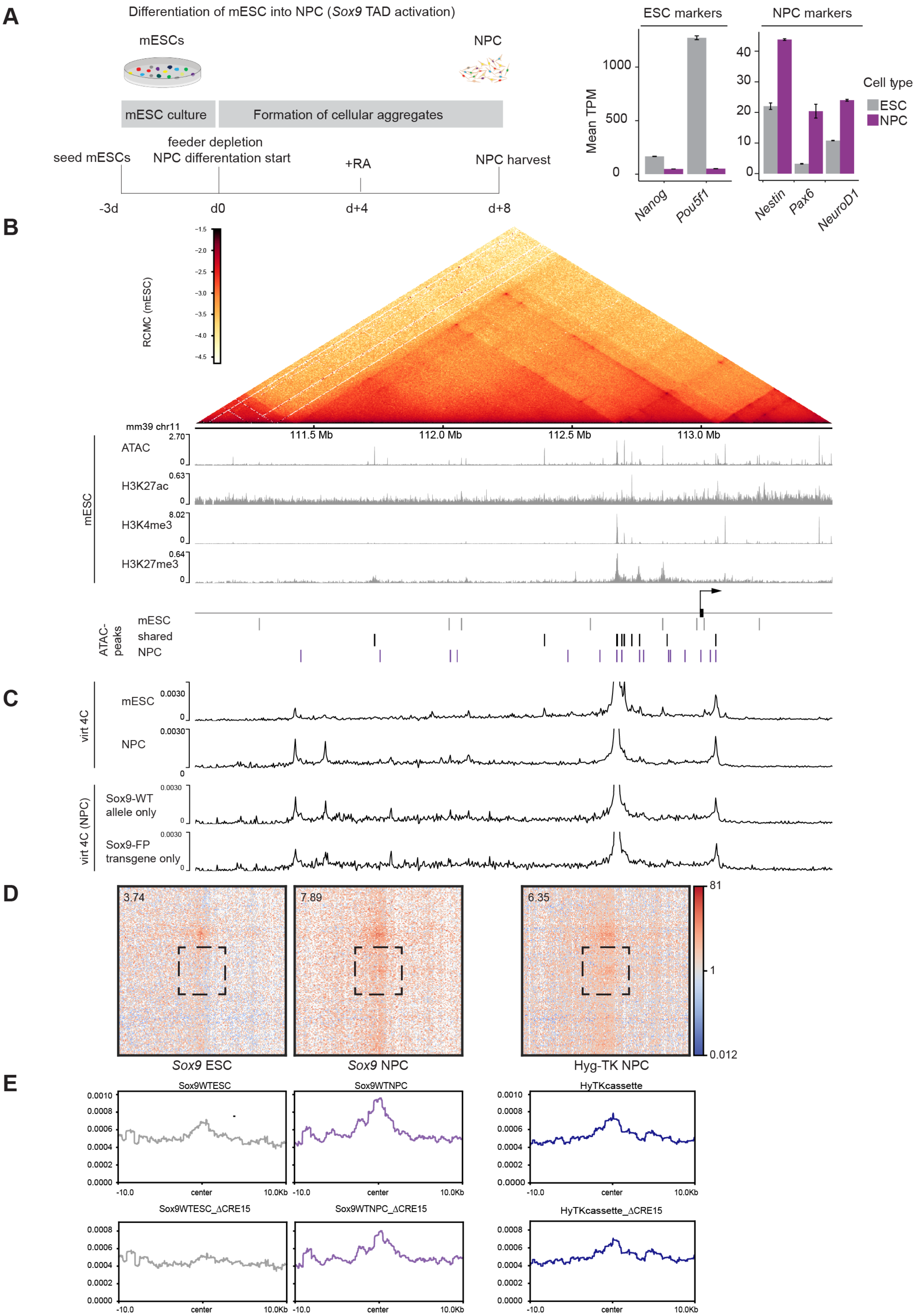


**Figure S1. NPC differentiation and Region-Capture Micro-C analysis**
(**A**) Schematic of NPC differentiation protocol (**left**) and switch in marker genes expression (RNA-seq) (**right**) (**B**) Region-Capture Micro-C map of the Sox9 locus in wildtype mESCs (top) and ATAC-seq/ChIP-seq tracks against histone modifications in mESC as well as mESC-specific (grey), shared (black) and NPC-specific ATAC-seq peaks (**C**) Virtual 4C tracks in mESC and NPCs from the Sox9 promoter (top) and in clonal cell lines with Sox9-FP transgene integrated, filtered for reads only mapping to the transgene (Sox9-FP) or endogenous Sox9 promoter. (**D**) 2D aggregate RCMC pileups of the Sox9 or Hyg-TK promoter (as indicated) with NPC-specific ATAC-seq peaks, values indicated normalized enrichment score. (**E**) 1D aggregate RCMC pileups as in (D), with and without CRE15, which is directly adjacent to a boundary CTCF site.

Supplemental Figure 1

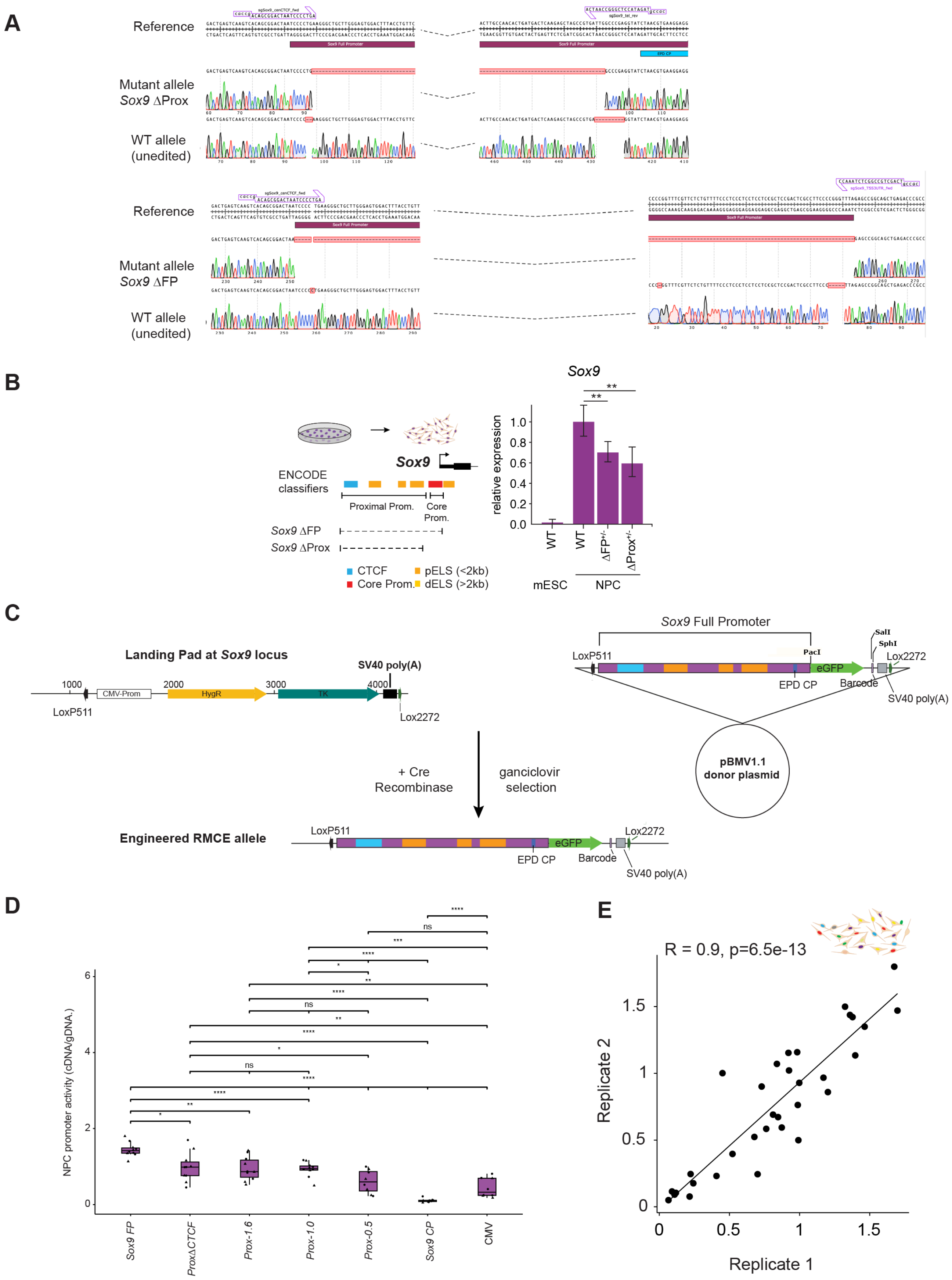


**Figure S2. Characterization of Sox9 promoter deletions, design of Promoter-RMCE targeting and measurements/reproducibility**
(**A**) Sanger sequencing traces of unmodified and genome edited alleles in mESC lines: ΔFP allele and ΔProx alleles. (**B**) Quantification of Sox9 expression in mESCs and NPCs in Sox9 promoter deletion cell lines using RT-qPCR (**C**) Schematic showing Hygromycin-Thymidin Kinase selection cassette integrated at the Sox9 locus, example donor plasmid carrying Sox9 Full Pomoter followed by eGFP and a barcode and its recombination replacing selection cassette (**D**) Results of Promoter-RMCE measurements with all significances reported p-values (* < 0.05, ** < 0.01, *** < 0.001, **** < 0,0001, n.s. = not significant (**E**) Reproducibility of Promoter-RMCE measurements at the Sox9 locus between two biological replicates of variant Sox9 promoter library in NPCs. R and p-values were calculated using the Pearson's correlation test.

**A**

| | Full promoter | | Core promoter | | | | | |
|---|---|---|---|---|---|---|---|---|
| **Gene** | **CpGi** | **CTCF** | **TATA** | **CCAAT** | **GC** | **INR** | **Full Promoter coordinates (mm10)** | **Length [bp]** |
| *Arhgap1* | + | - | - | - | + | - | chr2:91,649,252-91,650,307 | 1074 |
| *Snx3* | + | - | - | + | + | - | chr10:42,501,298-42,502,303 | 1036 |
| hPGK | + | - | - | + | + | - | chrX:78,103,817-78,104,327 (hg38) | 511 |
| *TDGF1* | - | - | - | - | + | - | chr9:110,946,019-110,948,275 | 2256 |
| *Sox2* | + | - | - | - | + | - | chr3:34,648,557-34,650,062 | 1505 |
| *Lefty1* | - | + | - | - | + | - | chr1:180,933,767-180,935,109 | 1342 |
| *Sox9* | + | + | - | + | + | - | chr11:112,779,569-112,782,402 | 2834 |
| *Pax6-1* | + | + | - | + | + | - | chr2:105,666,832-105,668,970 | 2138 |
| *Pax6-2* | + | - | - | + | + | - | chr2:105,673,488-105,675,719 | 2232 |
| *Vim* | - | - | + | - | + | + | chr2:13,572,238-13,574,369 | 2131 |
| *Nefm* | + | - | + | - | + | - | chr14:68,124,778-68,125,900 | 1122 |
| *Tbx5* | - | - | - | - | + | - | chr5:119,833,289-119,835,197 | 1910 |
| *Kcnj2* | - | + | - | - | + | - | chr11:111,064,703-111,066,247 | 1545 |
| *Kcnj16* | - | - | - | - | + | - | chr11:110,967,040-110,968,097 | 1058 |
| *Gata4* | + | - | - | + | + | - | chr14:63,245,159-63,247,300 | 2142 |
| *Eomes* | + | - | + | + | - | - | chr9:118,476,231-118,478,256 | 2026 |
| *Hey2* | + | - | - | + | + | - | chr10:30,842,716-30,844,900 | 2185 |
| *Fgfr2* | + | - | - | - | + | - | chr7:130,266,186-130,266,993 | 808 |

**B** Reproducibility of episomal MPRA (mESC) **C** Reproducibility of episomal MPRA (N2a)

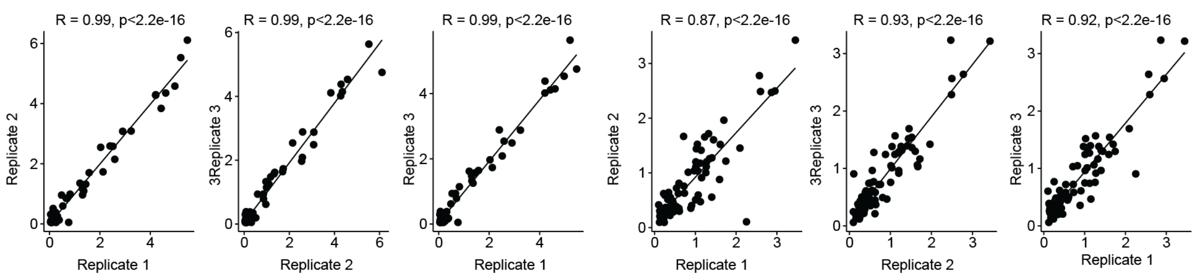


**D** Reproducibility of Promoter-RMCE (*β-globin*)

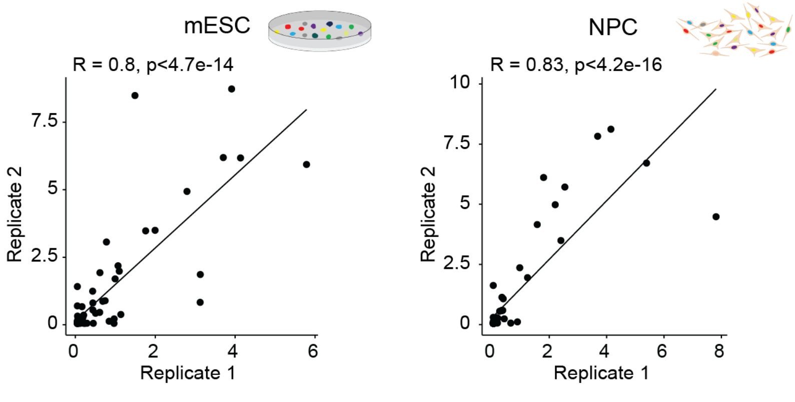


**Figure S3. Characterization of Full Promoter library. Reproducibility of episomal MPRA and Promoter-RMCE measurements at *β-globin***

(**A**) Sequence features of proximal promoter regions (annotated CpG islands, CTCF binding site) and known core promoter motifs (TATA-, CAAT- GC-Box, Initiator Motif) of promoters included in the promoter library, their genomic coordinates (mm10) and full promoter lengths (**B-C**) Reproducibility of full promoter episomal MPRA measurements from three biological replicates in mESCs (**B**) and N2a (**C**) cells. (**D**) Reproducibility of promoter-RMCE measurements at the β-globin locus between two biological replicates in mESCs (left) and NPCs (right). R and p-values were calculated using the Pearson's correlation test.

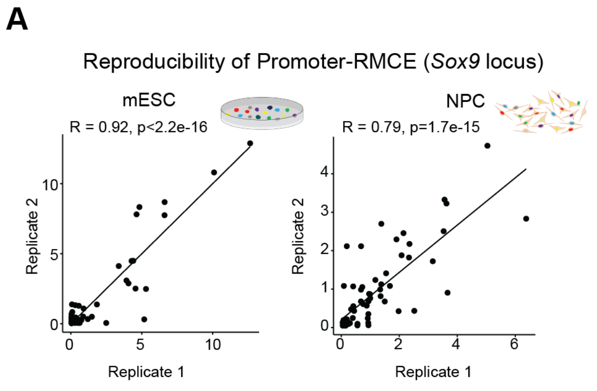


B

| Promoter | p_value | Median_activity | p_adj | fold_change_vs_CMV | log2FC vs CMV | Responder |
|---|---|---|---|---|---|---|
| hPGK | 1.554002e−04 | 3.54488169 | 0.0004403004 | 24.1131962 | 4.59175098 | TRUE |
| Arhgap1 | 4.570593e−05 | 2.32187440 | 0.0002590003 | 15.7939863 | 3.98130344 | TRUE |
| Snx3 | 1.371178e−03 | 0.93320297 | 0.0029137529 | 6.3478864 | 2.66627632 | TRUE |
| Lefty1 | 2.141414e−01 | 0.12371830 | 0.2800310800 | 0.8415636 | −0.24885573 | FALSE |
| TDGF1 | 3.791764e−02 | 0.07733936 | 0.0585999859 | 0.5260822 | −0.92663978 | FALSE |
| Sox2 | 5.737374e−01 | 0.15716022 | 0.6095959596 | 1.0690442 | 0.09632149 | FALSE |
| Sox9 | 4.570593e−05 | 1.68332757 | 0.0002590003 | 11.4504267 | 3.51732946 | TRUE |
| Nefm | 6.660007e−04 | 1.01594949 | 0.0016174302 | 6.9107495 | 2.78884218 | TRUE |
| Vim | 2.930403e−02 | 1.00930238 | 0.0498168498 | 6.8655342 | 2.77937197 | TRUE |
| Gata4 | 4.444444e−02 | 0.87959847 | 0.0629629630 | 5.9832548 | 2.58093050 | FALSE |
| Fgfr2 | 1.554002e−04 | 0.87219304 | 0.0004403004 | 5.9328812 | 2.56873289 | TRUE |
| Kcnj2 | 4.570593e−05 | 0.81546785 | 0.0002590003 | 5.5470219 | 2.47171342 | TRUE |
| Hey2 | 1.554002e−04 | 0.45931815 | 0.0004403004 | 3.1244001 | 1.64357921 | TRUE |
| Eomes | 4.387769e−03 | 0.24514127 | 0.0082880083 | 1.6675139 | 0.73769882 | TRUE |
| Pax6_1 | 5.696970e−01 | 0.10288190 | 0.6095959596 | 0.6998291 | −0.51492534 | FALSE |
| Kcnj16 | 5.726039e−01 | 0.09894084 | 0.6095959596 | 0.6730210 | −0.57127654 | FALSE |
| Tbx5 | 6.620047e−01 | 0.09258057 | 0.6620046620 | 0.6297568 | −0.66713324 | FALSE |

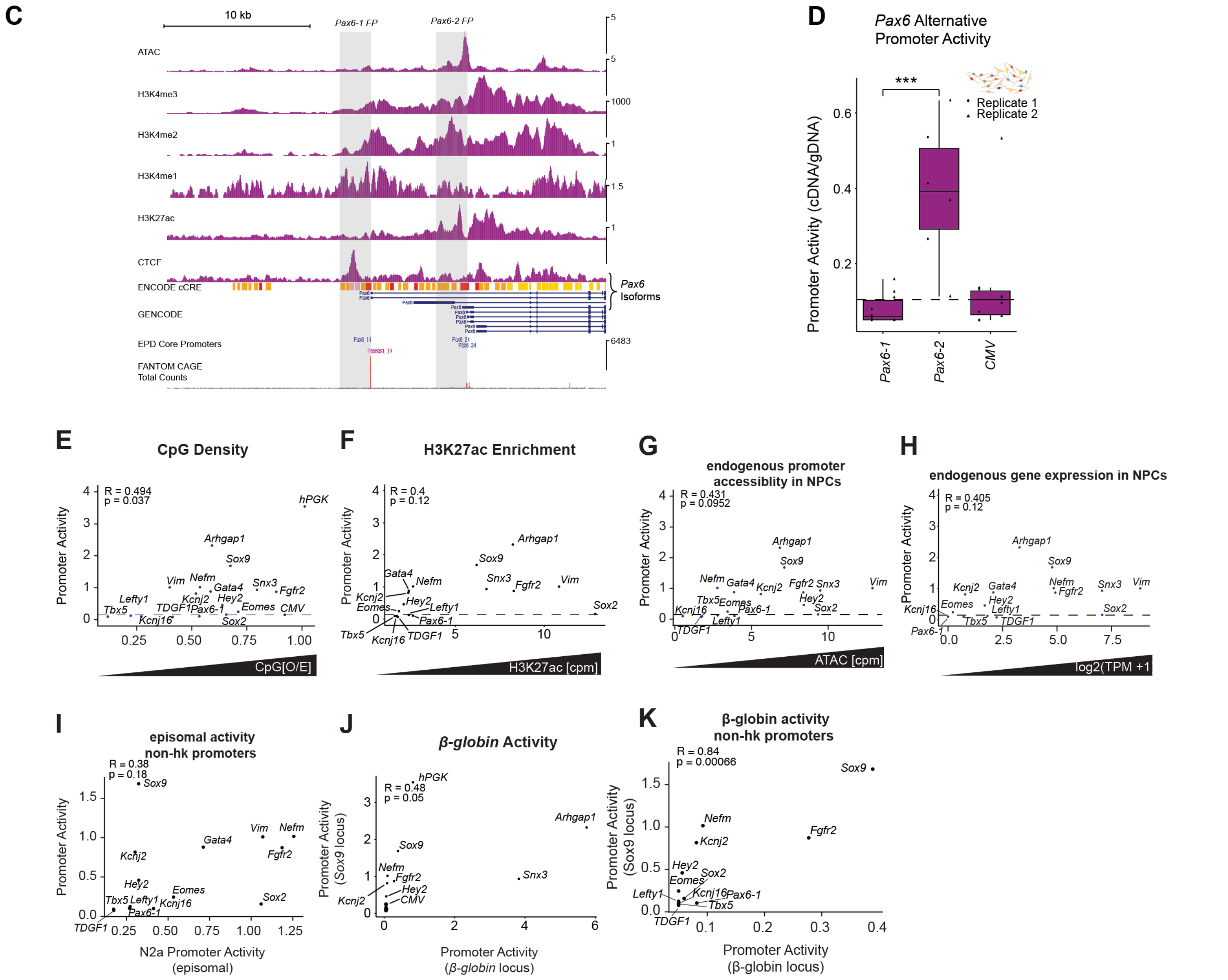


**Figure S4. Reproducibility of Promoter-RMCE measurements at the Sox9 locus, activity of alternative Pax6 promoters and correlation between activity of full promoters and endogenous gene expression, promoter accessibility or episomal activity**

(**A**) Promoter-RMCE reproduciblity between two biological replicates (full promoter library at Sox9 locus) in mESCs (left) and NPCs (right). (**B**) Statistical analysis of promoter activity relative to CMV control; Two-sided Wilcoxon rank-sum test at barcode resolution. p-values were adjusted for multiple testing using the Benjamini–Hochberg procedure. Median fold-changes relative to CMV and responder classification (BH-adj. p-value < 0.05 and FC > 1.5).

(**C**) NPC chromatin marks, ENCODE cCRE annotation and FANTOM Total CAGE counts at the Pax6 locus (**D**) NPC Promoter-RMCE activity at Sox9 locus shows differential response of Pax6-1 and Pax6-2 promoters. Triangles and dots = barcode values from two biological replicates. Dashed line = median CMV activity. p-value < 0.05 from Wilcoxon test. (**E-H**) Median promoter activity at the Sox9 locus in NPCs as a function of promoter CpG density (E) H3K27ac signal at the endogenous promoter (F) endogenous promoter accessibility (ATAC-seq CPM in the TSS −3 kb/+2 kb window) in NPCs or endogenous gene expression (RNA-seq log2 TPM) in NPCs (H). (**I**) Median promoter activity at the Sox9 locus in NPCs as a function of promoter activity in N2a cells episomally (**I**) or genome-integrated at the β-globin locus for for all (**J**) or only non-housekeeping full promoters (**K**). Pearson's correlation coefficient (R) and P values were calculated using Pearson's correlation test.

Supplemental Figure 4

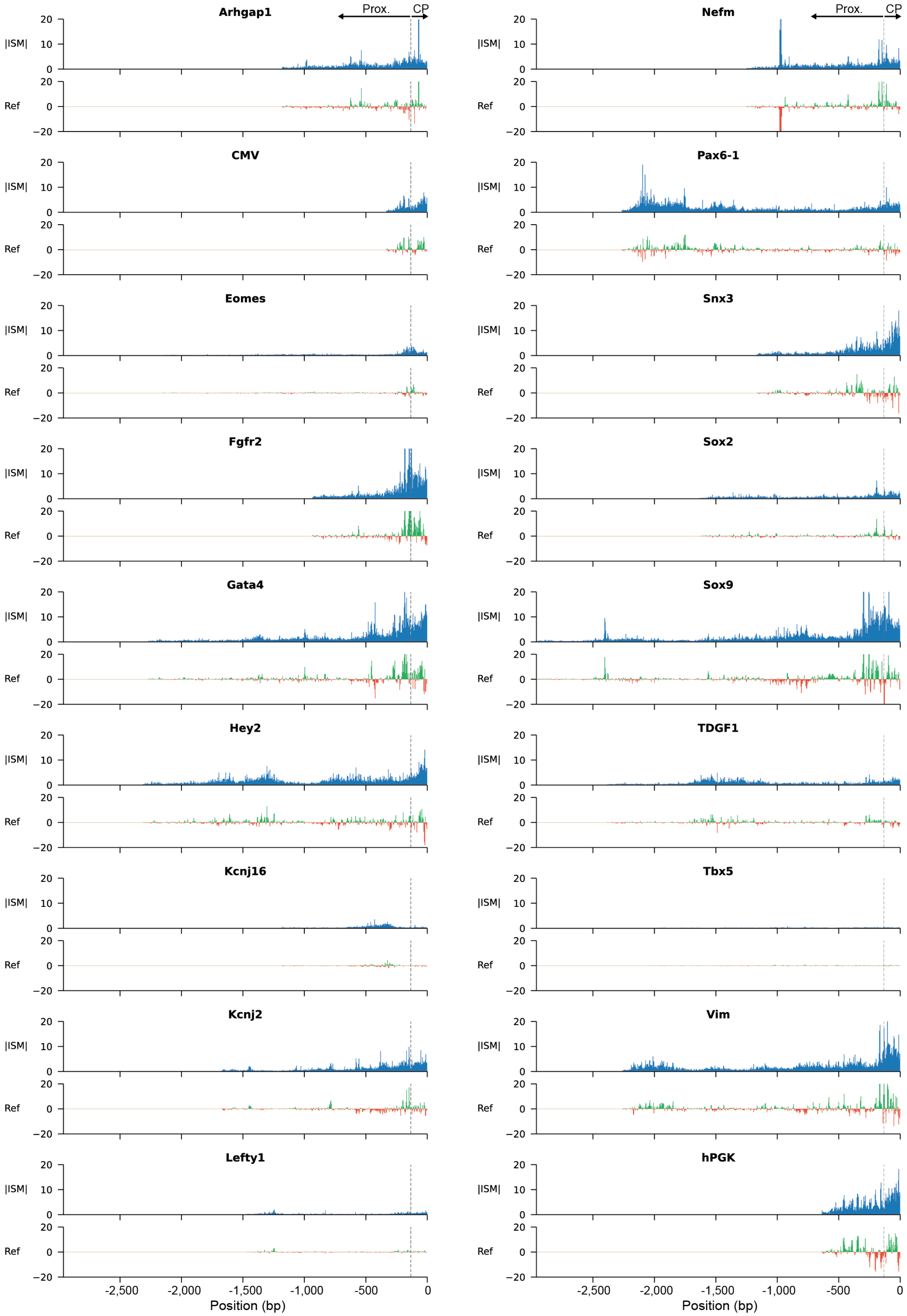


**Figure S5. ISM-based nucleotide contribution scores for all full promoters in the library.**

(Top) Absolute ISM-based contribution scores, calculated by adding up the absolute value of nucleotide contributions for all possible alleles at each position (Bottom) Contribution scores for the reference construct sequence. Green values indicate positive, and red values indicate negative contribution. Dashed line indicates separation of annotated core promoter and proximal promoter region

Supplementary Figure 5

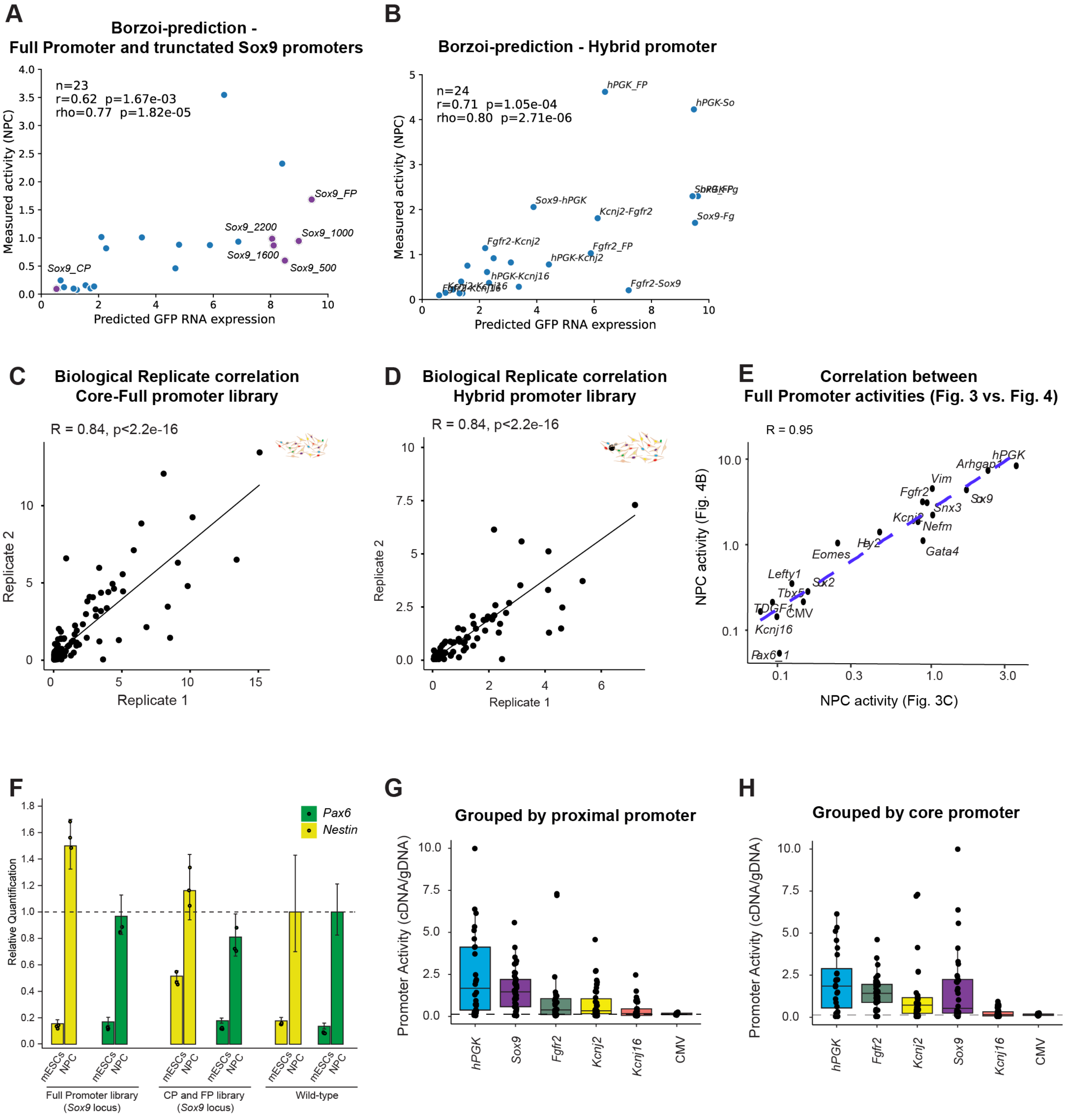


**Figure S6. Reproducibility of Promoter-RMCE measurements for core vs. full promoter and hybrid promoter experiments**
**(A)** Measured Promoter activity in NPCs of Full and truncated Sox9 promoters as a function Borzoi-ML model predicted activity. Truncated Sox9 Promoters are highlighted in purple (B) Measured Promoter activity in NPCs of Hybrid promoter library as a function Borzoi-ML model predicted activity. (**C**) Reproducibility of Promoter-RMCE measurements at the Sox9 locus between two biological replicates of core and full promoter library in NPCs. (**D**) Reproducibility of Promoter-RMCE measurements at the Sox9 locus between two biological replicates of hybrid library in NPCs.(**E**) Correlation between Full Promoter activity in NPCs as measured for Fig. 3C (x-axis) and Fig. 4 (y-axis). Note reproducibility despite different scales for cDNA/gDNA values, due to amplification bias in Amplicon-seq (**F**) Relative expression of Nestin and Pax6-2 in mESC and NPC samples of selected RMCE experiments measured by RT-qPCR, normalized to Gapdh and quantified using the 2^-ΔΔCt method. Wildtype NPCs were used as calibrator sample. Bars show mean realtive expression, error bars represent the ΔΔCt-derived confidence interavl. Individual datapoints correspond to technical qPCR replicates (n=3). (**G, H**) Boxplots depict median response rate of hybrid promoters grouped by proximal and core promoters. Dashed line: median CMV activity.
